# Promoting Fc-Fc interactions between anti-capsular antibodies provides strong immune protection against *Streptococcus pneumoniae*

**DOI:** 10.1101/2022.01.21.477211

**Authors:** Leire Aguinagalde, Maurits A. den Boer, Suzanne M. Castenmiller, Seline A. Zwarthoff, Carla J.C Gosselaar-de Haas, Piet C. Aerts, Frank J. Beurskens, Janine Schuurman, Albert J.R. Heck, Kok P.M. van Kessel, Suzan H.M. Rooijakkers

## Abstract

*Streptococcus pneumoniae* is the leading cause of community-acquired pneumonia and an important cause of childhood mortality. Despite the introduction of successful vaccines, the global spread of both non-vaccine serotypes and antibiotic-resistant strains reinforce the development of alternative therapies against this pathogen. One possible route is the development of monoclonal antibodies (mAbs) that induce killing of bacteria via the immune system. Here we investigate whether mAbs can be used to induce killing of pneumococcal serotypes for which the current vaccines show unsuccessful protection. Our study demonstrates that when human mAbs against pneumococcal capsule polysaccharides (CPS) have a poor capacity to induce complement activation, a critical process for immune protection against pneumococci, their activity can be strongly improved by hexamerization-enhancing mutations. Our data indicate that anti-capsular antibodies may have a low capacity to form higher-order oligomers (IgG hexamers) that are needed to recruit complement component C1. Indeed, specific point mutations in the IgG-Fc domain that strengthen hexamerization strongly enhance C1 recruitment and downstream complement activation on encapsulated pneumococci. Specifically, hexamerization-enhancing mutations E430G or E345K in CPS6-IgG strongly potentiate complement activation on *S. pneumoniae* strains that express capsular serotype 6 (CPS6), and the highly invasive serotype 19A strain. Furthermore, these mutations improve complement activation via mAbs recognizing CPS3 and CPS8 strains. Importantly, hexamer-enhancing mutations enable mAbs to induce strong phagocytosis and intracellular killing by human neutrophils. Finally, passive immunization with CPS6-IgG1-E345K protected mice from developing severe pneumonia. Altogether, this work provides an important proof-of-concept for future optimization of antibody therapies against encapsulated bacteria.

## Introduction

The Gram-positive bacterium *Streptococcus pneumoniae* (pneumococcus) is the leading cause of community-acquired pneumonia and a major cause of bacteraemia and meningitis in children and adults (1–3). While pneumococcus commonly resides asymptomatically in the nasopharynx, it can cause a wide spectrum of infections in children, elderly and immunocompromised patients (4–7). Infections by pneumococcus range from non-invasive diseases, such as otitis media and sinusitis to life-threatening bacteremia and meningitis. To reduce its great impact on morbidity and mortality, vaccines have been successfully developed and introduced world-wide. Currently available vaccines target the polysaccharide capsule (CPS), which is considered the most important virulence factor of *S. pneumoniae*. Although there are more than 90 different capsular serotypes (8), the current vaccines only include a limited number of serotypes including those most frequently found to be causing invasive pneumococcal disease (IPD). Besides the fact that the widespread vaccination has been highly effective in lowering IPD caused by vaccine serotypes (9–11), there still is a large burden of pneumococcal disease caused by non-vaccine serotypes. Furthermore, because some vaccine serotypes induce a weak immune response it remains difficult to control pneumococcal disease, particularly in risk groups (12). Finally, the emergence of strains with high level of antibiotic resistance (13–16) highlight a strong need to develop new therapeutic strategies against pneumococcal infections.

In recent years, antibody therapies have emerged as a successful treatment for several autoimmune diseases and cancers (17, 18). Therefore, there is now also great interest in development of antibody-based therapies against bacterial infections. To eliminate bacteria, antibodies should bind to the bacterial surface and induce killing via the immune system. As evidenced by recurrent infections in patients with genetic complement deficiencies (19, 20), human immune protection against pneumococci critically depends on the action of the human complement system (21, 22). Complement is a large network of proteins in blood and other body fluids. These proteins circulate as inactive precursors but become rapidly activated upon contact with bacterial cells (23). An activated complement cascade triggers a variety of immune responses, such as the labeling of bacteria with C3-derived opsonins (C3b and iC3b) that potently induce phagocytosis and intracellular killing of bacteria by professional phagocytes (24, 25).

Because complement is essential in immune protection against *S. pneumoniae* (26–28), the capacity of antibodies to induce complement activation could be exploited for effective antibacterial therapies. However, it is not known whether monoclonal antibodies effectively induce complement activation on *S. pneumoniae.* This is especially unclear for encapsulated *S. pneumoniae* strains, because the capsule is believed to block immune activation, for instance by shielding epitopes and blocking deposition of C3 opsonins (29). Furthermore, recent studies showed that target-bound IgGs should organize into higher-order oligomers (IgG hexamers) to provide an optimal docking platform for complement component C1q (30, 31) (**Figure 1a and S1**). Because each antibody-binding headpiece of C1q has a low affinity for IgG, physiological binding occurs when the six headpieces simultaneously bind to IgG once hexamerized. Here we study the potential of several anti-capsular monoclonal antibodies to induce complement activation and phagocytic killing of encapsulated pneumococci. Our data suggest that these anti-capsular antibodies as wild-type IgG have a poor capacity to form IgG hexamers and trigger downstream killing via complement. Importantly, this limitation can be overcome by introduction of a single amino acid mutation that enhances hexamerization of anti-capsular antibodies and strongly potentiates complement-mediated killing of pneumococci, both *in vitro* and *in vivo*.

**Figure 1.**
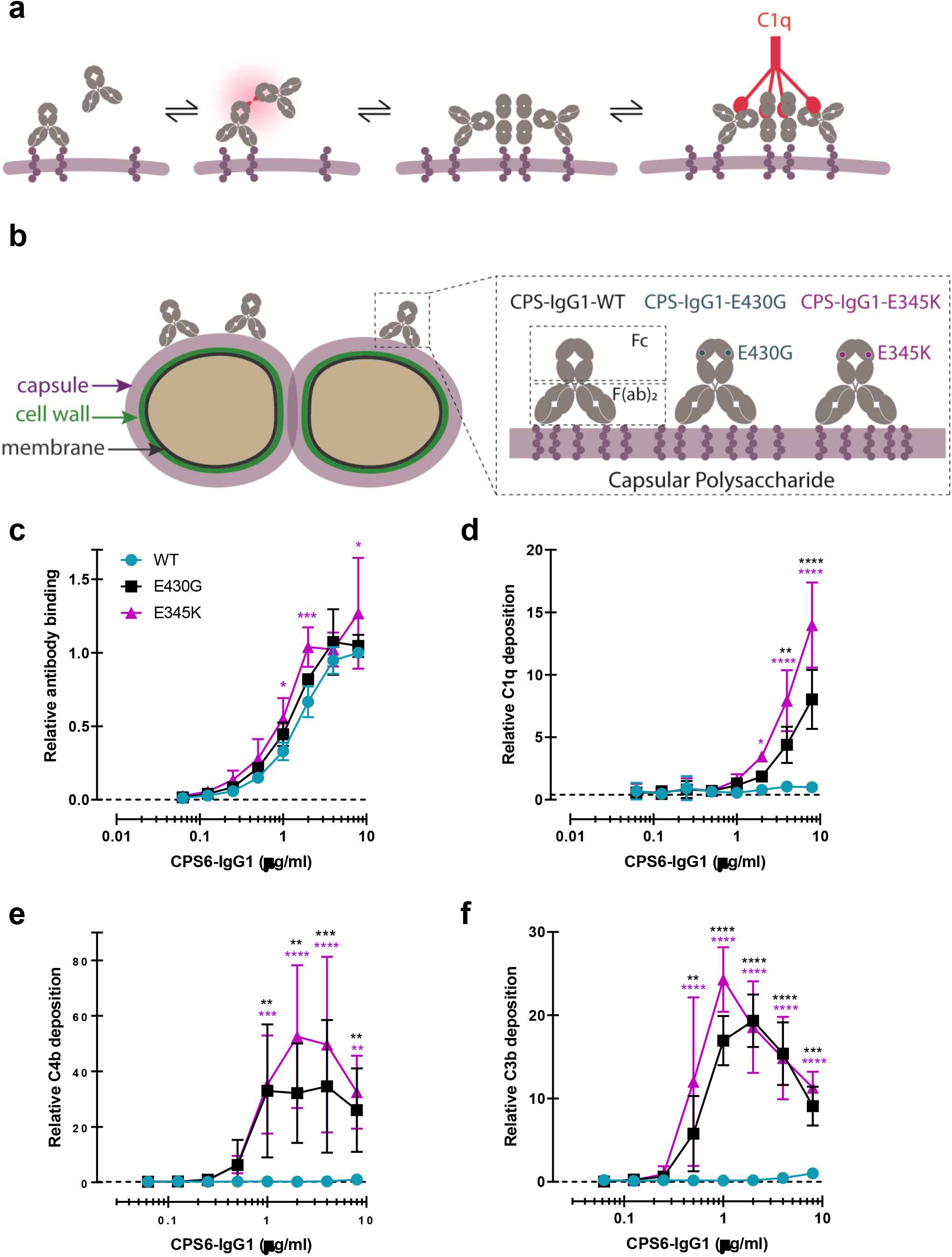
Promoting Fc-Fc interactions between CPS6-IgG1 enables complement activation on *S. pneumoniae* 6B. (a) Schematic representation of antibody binding to antigen on a target surface. IgGs can cluster into hexamers via non-covalent interaction between their Fc domains and thus form an optimal docking platform for C1q. (b) *Left*, schematic illustration of *S. pneumoniae* showing the location of its dominant surface structure, the polysaccharide capsule (CPS), and antibodies recognition that confer type-specific protection. The capsule forms the outermost layer of encapsulated strains of *S. pneumoniae* and for most cases is covalently attached to the outer surface of the cell wall peptidoglycan. *Right,* binding of CPS-IgG1wild-type (WT) or containing the single point hexamerization-enhancing mutations, E430G or E345K, to *S. pneumoniae* surface. (c) Binding of WT and hexamerization-enhancing mutated (E430G or E345K) CPS6-IgG1 to *S. pneumoniae* 6B (ST6B), detected with Alexa^647^-conjugated F(ab’)2-goat anti-human Kappa antibody by flow cytometry. (d-f) Complement components C1q, C4b and C3b deposition on *S. pneumoniae* 6B after incubation with 2.5% IgG/IgM-depleted serum supplemented with WT or hexamerization-enhancing mutated (E430G or E345K) CPS6-IgG1. All detected with Alexa^647^-conjugated F(ab’)2-goat anti-mouse immunoglobulins antibody by flow cytometry. (c-f) Data are expressed relative to the 8 µg WT value and presented as means ± SD of three independent experiments. Dashed line represents background (no IgG) level. Two-way ANOVA was used to compare across dose-response curves at the various concentrations the differences between the WT and the E430G or E345K variants. When significant it was displayed as *P < 0.05; ***P < 0.001; ****P < 0.0001.

## Results

### Hexamer-enhancing mutations enable anti-capsular antibodies to activate complement on *S. pneumoniae* serotype 6B

To study whether human mAbs can induce complement activation on encapsulated *S. pneumoniae* strains, we first investigated complement activation on *S. pneumoniae* 6B, a common serotype infecting both adults and children (32, 33). Although serotype 6B is covered by current vaccines, the 6B capsule type is found to be poorly immunogenic and an important risk factor in the mortality by IPD (32–34). We generated recombinant variants of a previously identified human IgG1 antibody that recognizes a carbohydrate structure present on serogroup 6 strains: α-D-Glcp(1→3)α-L-Rhap (35). The variable, antigen-binding (Fab) domain of this polysaccharide serogroup 6-specific antibody (CPS6-IgG or ‘Dob1’(35)) was cloned into expression vectors containing the constant (Fc) domains of human IgG1. Furthermore, we introduced single amino acid mutations in the IgG1 Fc domain to enhance Fc-dependent hexamerization of target-bound antibodies (**Figure 1b**) (23,30,31). Specifically, mutations E430G (Glu^430^ → Gly) or E345K (Glu^345^ →Lys) were introduced because of their proven strong enhancement of complement-dependent lysis of tumour cells while retaining properties required for the development of biopharmaceuticals (30, 31).

After verifying that the introduction of hexamer-enhancing mutations did not affect the binding of CPS6-IgG1 antibodies to serotype 6B (**Figure 1c and S2**), we studied their capacity to induce complement activation. To this end, 6B pneumococcus was incubated with mAbs in the presence of human serum as complement source. To exclude the involvement of pre-existing antibodies, we used human serum that was depleted from naturally occurring IgG and IgM (36) (denoted IgG/IgM-depleted serum). Using surface-specific staining of C1q and flow cytometry, we first determined the capacity of mAbs to recruit C1q (Figure S1). While the wild-type (WT) CPS6-IgG1 showed little to no reactivity with C1q, we noted that the introduction of hexamer-enhancing mutations E430G or E345K strongly enhanced the ability to interact with C1q (**Figure 1d)**. Importantly, we found that introduction of hexamer-enhancing mutations in CPS6-IgG1 allowed activation of the classical complement pathway. Recruitment of C1q to target-bound IgGs induces activation of C1q-attached C1r and C1s proteases that cleave C4 to covalently attach activated C4b molecules onto the bacterial surface (**Figure S1**) (37). Furthermore, C1 activates C2 to produce a C3 convertase (C4b2b) that deposits large amounts of C3b, a key component of the complement cascade that labels bacteria for phagocytosis (38, 39). Indeed, by monitoring deposition of C4b and C3b molecules, we observed that Fc-engineered variants of CPS6-IgG1, but not the WT antibody, potently induced deposition of C4b (**Figure 1e)** and C3b molecules (**Figure 1f)** onto serotype 6B. Altogether, these data show that hexamer-enhancing mutations can overcome poor complement activation by monoclonal antibodies against capsular serotype 6.

### Hexamer-enhancing mutations in CPS6-IgG enhance phagocytosis of *S. pneumoniae* serotype 6B

Next, we investigated whether the enhanced complement activation also impacted phagocytosis of serotype 6B by human neutrophils. Neutrophils are crucial to establish immune protection against pneumococcal infections (40, 41). These cells are the first to be recruited from the blood to the site of infection where they engulf and internalize bacteria via phagocytosis to, subsequently, kill them by exposure to antimicrobial agents such as antimicrobial peptides, reactive oxygen species and enzymes (22). The phagocytic uptake is greatly enhanced by the tagging of bacteria with IgG antibodies that can engage Fcγ receptors (FcγRs) (42) and/or C3-derived opsonins that can mediate the uptake of bacteria via complement receptors (25, 43). We studied phagocytosis of fluorescent serotype 6B by freshly isolated human neutrophils (44) in the presence of CPS6-IgG1 mAbs and IgG/IgM-depleted human serum as complement source. First, we observed that the CPS6-IgG1-WT antibody poorly induced phagocytosis of serotype 6B (**Figure 2a and S3**). In contrast, hexamer-enhanced variants of CPS6-IgG1 induced very potent phagocytosis (**Figure 2a and S3a**). For both E430G and E345K mutated variants, we found that 0.3 μg/ml of mAb induced maximum phagocytosis in the presence of 2.5% IgG/M-depleted serum (**Figure 2a**). When serum was heat-treated to inactivate complement (45), we found that phagocytosis by E430G and E345K antibodies was completely abolished (**Figure 2b)**. Bacterial internalization in presence of active complement and antibodies was confirmed by microscopy (**Figure 2c**). This indicates that phagocytic uptake via hexamerization-enhanced CPS6 antibodies is fully depended on the presence of complement.

**Figure 2.**
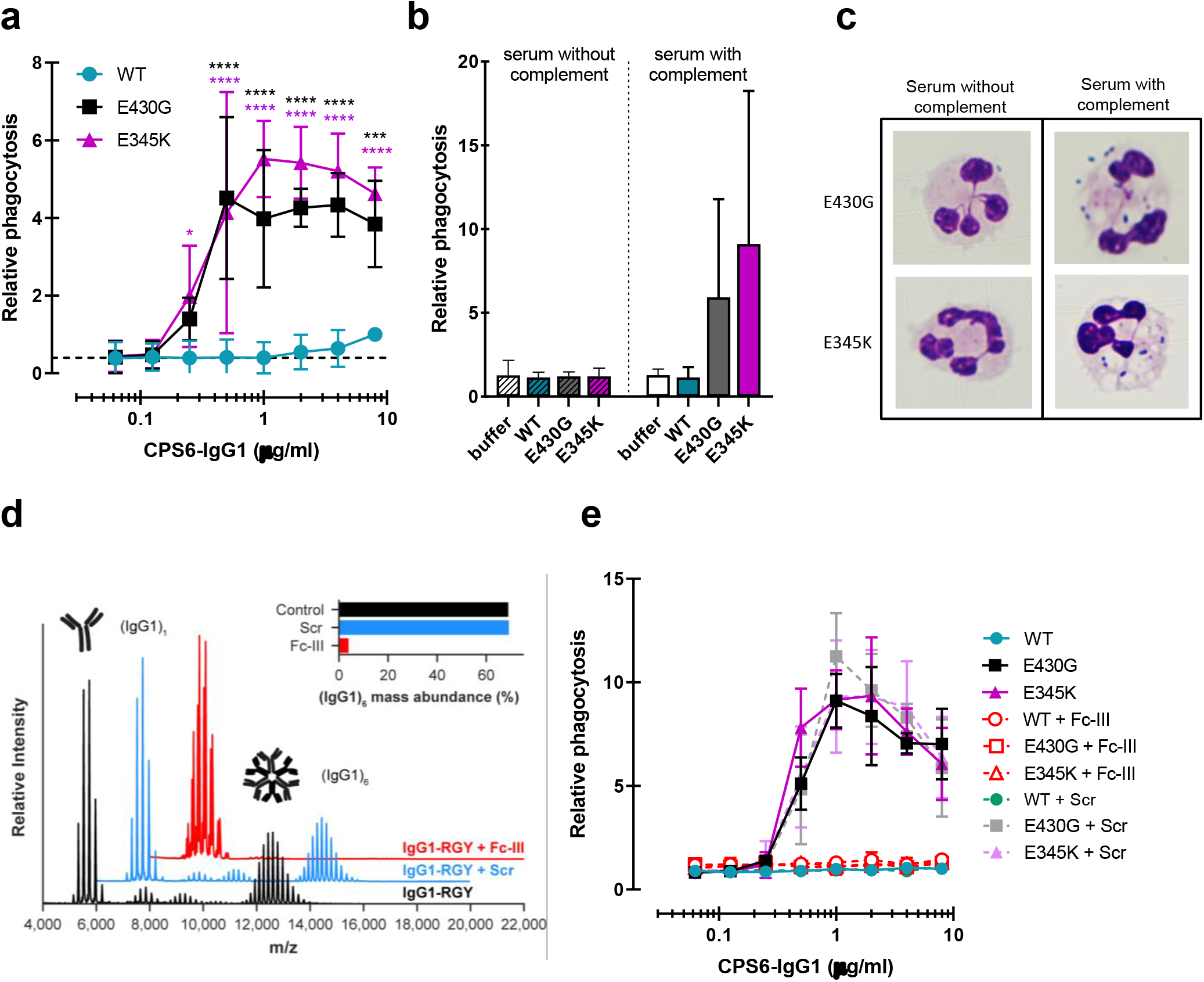
Hexamerization-enhanced variants of CPS6-IgG1 trigger complement-dependent phagocytosis of serotype 6B *S. pneumoniae*. (a) Phagocytosis in the presence of complement. Fluorescently labelled bacteria uptake by human neutrophils in the presence of 2.5% IgG/IgM-depleted serum supplemented with CPS6-IgG1-WT versus E430G and E345K variants. (b) Comparison of *S. pneumoniae* serotype 6B phagocytosis by CPS6-IgG1-WT, E430G or E345K antibody variants at 4 µg/ml in the presence of 2.5% IgG/IgM-depleted serum without (striped bars) or with active complement cascade (non-striped bars). (c) Microscopy image of pneumococcal phagocytosis by human neutrophils in the presence of 5% IgG/IgM-depleted serum with or without active complement, supplemented with 8 µg/ml CPS6-IgG1-E430G or E345K variant. Cytospin preparations were stained with Giemsa–May–Grünwald (Diff-Quik) and pictures taken using a 100X objective to visualize cytoplasmic internalization. (d) Native mass spectra of IgG1-RGY in the absence (black) and presence of Fc-Fc inhibitor peptide Fc-III (red) or a scrambled version Scr (blue). Spectra are shifted for clarity and monomeric (IgG1)_1_ and hexameric (IgG1)_6_ mass peaks are indicated. Inset represents the percentage (IgG1)_6_ for each sample. (e) Phagocytosis of fluorescently labelled *S. pneumoniae* 6B after incubation with 2.5% IgG/M-depleted serum supplemented with CPS6-IgG1-WT, CPS6-IgG1-E430G or CPS6-IgG1-E345K, in presence or absence of 10 μg/ml Fc-Fc inhibitory peptide (Fc-III) and a scrambled version (Scr). (a, b, e) Bacterial uptake is displayed as the mean fluorescence value of neutrophils relative to CPS6-IgG1-WT at the highest concentration tested (8 μg/ml). Data represent mean ± SD of three independent experiment. (a) Dashed line represents background (buffer) level. Two-way ANOVA was used to compare across dose-response curves at the various concentrations the differences between the WT and the E430G or E345K variants. When significant it was displayed as *P < 0.05; ***P < 0.001; ****P < 0.0001.

To study whether phagocytosis via hexamer-enhanced antibodies indeed depends on Fc-dependent IgG oligomerization, we analyzed phagocytosis in the presence of Fc-III, a cyclic peptide that binds to IgG residues involved in the Fc-Fc interaction interface (46). To study this end we combined native mass spectrometry (47, 48) with an IgG triple mutant (IgG-RGY, combination of E345R (Glu^345^ → Arg), E430G (Glu^430^ → Gly) and S440Y (Ser^440^ → Tyr) mutations) that has the capacity to form stable hexamers in solution (30). In agreement with prior work, native mass spectra of IgG1-RGY revealed the presence of monomeric and hexameric species, with intermediate states observed at lower abundance (**Figure 2d**) (23, 30). When IgG1-RGY was incubated with Fc-III however, the relative abundance of IgG oligomers was diminished. No effect was observed for a scrambled version of the same peptide (Scr) that in contrast to Fc-III does not bind IgG1 molecules (**Figure 2d and S4**). The fact that we observed binding of Fc-III to monomeric IgG1, but not to larger oligomeric species, suggests that Fc-III inhibits IgG-mediated complement activation by competitive binding to the Fc-Fc interaction interface of IgG monomers. In line with our hypothesis, we found that Fc-III potently blocked phagocytic uptake of serotype 6B via both E430G and E345K mutants, while Scr showed no effect (**Figure 2e**).

Taken together, while a WT monoclonal antibody against CPS6 has a poor capacity to induce phagocytosis of *S. pneumoniae* serotype 6B, introduction of hexamerization-enhancing mutations strongly increases complement-mediated phagocytosis.

### Introduction of hexamer-enhancing mutations in IgG2 and IgG3 also results in increased pneumococcal recognition and clearance

Because the natural antibody response against bacterial capsule polysaccharides is dominated by IgG2 (49–51), we also constructed human monoclonal CPS6-IgG2. While WT and Fc-Fc enhancing variants showed equal binding to serotype 6B (**Figure S5a**), we again observed that hexamer-enhancing mutations E430G and E345K both improved C3b deposition in IgG2 (**Figure 3a**).Also for IgG3 antibodies, which are considered more effective in the induction of Fc-effector functions (52), we found that hexamer-enhancing mutation E345K significantly increased C3b deposition, while a much less strong enhancement was observed for E430G (**Figure 3b**). A direct comparison of E345K variants shows that CPS6-IgG1-E345K activated complement more potently than CPS6-IgG3-E345K and CPS6-IgG2-E345K (**Figure S5e),** even though the binding is the same (**Figure S5c**).

**Figure 3.**
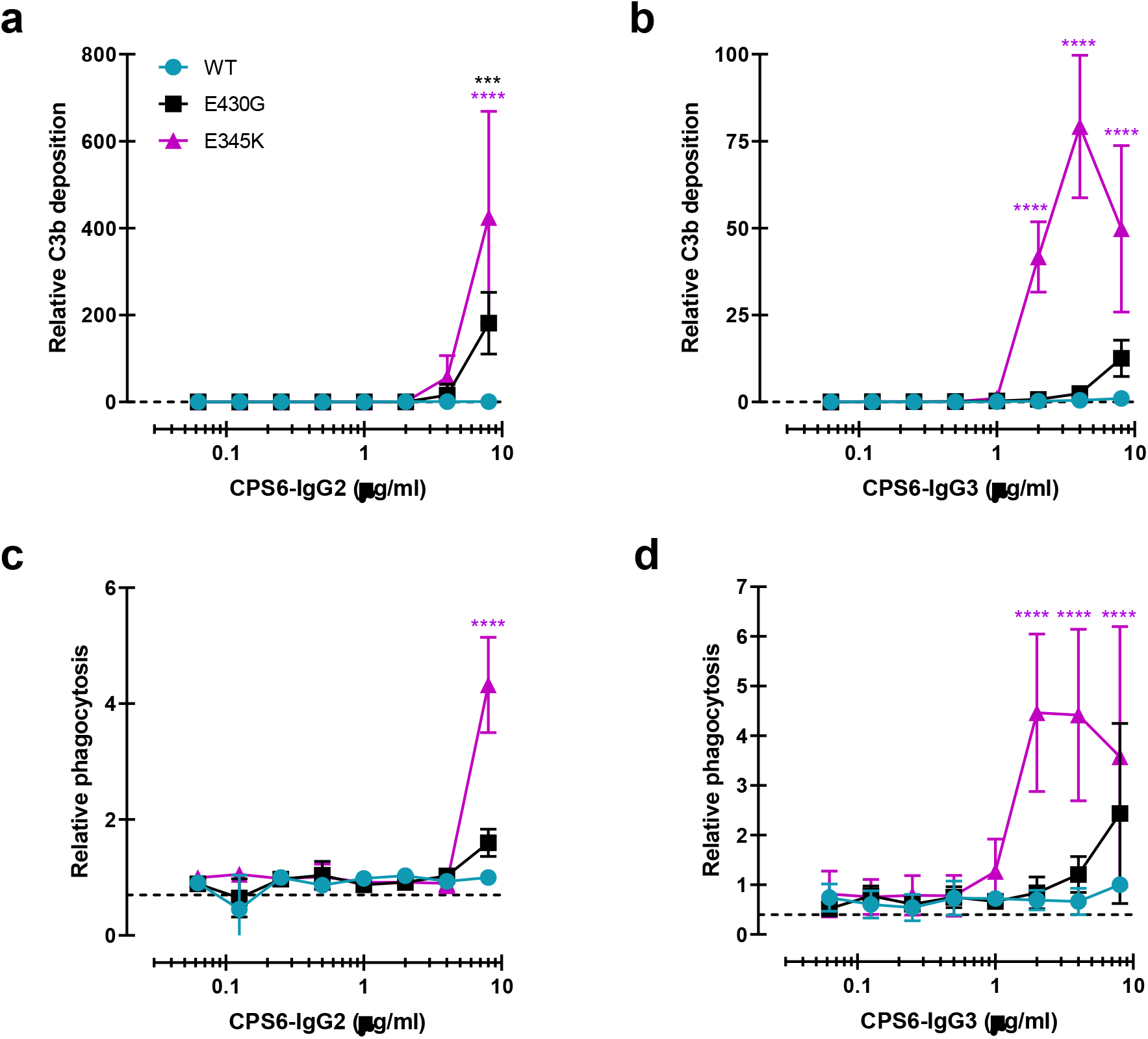
Introduction of E430G or E345K mutation in CPS6-IgG2 and CPS6-IgG3 improves complement activation and phagocytosis of *S. pneumoniae*. (a, b) C3b deposition on serotype 6B surface after incubation of bacteria with CPS6-IgG2 (a) or CPS6-IgG3 (b) antibody variants, in presence of 2.5% IgG/IgM-depleted serum and detected with a monoclonal murine anti-human C3d antibody by flow cytometry. (c, d) Fluorescent serotype 6B bacterial phagocytosis by human neutrophils detected by flow cytometry after incubation in the presence of CPS6-IgG2 (c) or CPS6-IgG3 (d) hexamerization-enhanced variants, E430G and E345K, plus 2.5% IgG/IgM-depleted serum. All data are presented as mean fluorescence relative to the highest CPS-IgG-WT concentration tested (8 µg/ml). Dashed line represents background (no IgG) level. Data represent mean ± SD of at least two independent experiments. Two-way ANOVA was used to compare across dose-response curves at the various concentrations the differences between the WT and the E430G or E345K variants. When significant it was displayed as *P < 0.05; ***P < 0.001; ****P < 0.0001.

Consistent with the results for C3b deposition, we observed that E345K, but not E430G, enhanced phagocytosis of CPS6-IgG3 (**Figure 3d**) while a very moderate effect was observed for IgG2 (**Figure 3c**). Again, phagocytosis of CPS6-IgG3 fully relied on the presence of active complement (**Figure S5f**).

### CPS6-IgG1-E430G and CPS6-IgG1-E345K induce potent complement activation and phagocytosis of serotypes 6A, 6C and 19A

We wondered whether these results could be extended to other pneumococcal serotypes. Next to serotype 6B, the α-D-Glcp(1→3)α-L-Rhap antigen is also found in the CPS of serotypes 6A, 6C and 19A but not in the 19F CPS (35). Serotype 19A is of particular interest because this is a highly invasive serotype that, despite coverage in PCV-13 vaccine, remains one of the most frequently carried pneumococcal serotypes in children, and major cause of disease in European countries and the USA (53–56). Furthermore, there is a significant increase in penicillin and multidrug resistance among 19A clinical isolates (57–60). After validating that (hexamer-enhancing variants of) CPS6-IgG1 indeed bind to serotype 19A but not to 19F (**Figure S6a,d**), we tested complement activation and phagocytosis of this strain. Similar to our results on serotype 6B, we observed that hexamer-enhancing mutations strongly increased complement activation on serotype 19A, as evidenced by increased detection of C1q (**Figure 4a**), C4b (**Figure 4b**) and C3b (**Figure 4c**), especially when E345K mutation was present. Consistently, CPS6-IgG1-E430G and CPS6-IgG1-E345K enhanced phagocytosis of *S. pneumoniae* serotype 19A phagocytosis in a dose-dependent manner **(Figure 4d**). Finally, we validated that hexamer-enhancing variants of CPS6-IgG1 also bind and improve complement-dependent phagocytosis of the poorly immunogenic serotype 6A (61) **(Figure 4e and S5b**) and the non-vaccine serotype 6C **(Figure 4f and S6c).** Altogether these results suggest that hexamer-enhancing variants of CPS6-IgG1 trigger complement-dependent phagocytosis of serogroup 6 pneumococci and the highly invasive serotype 19A.

**Figure 4.**
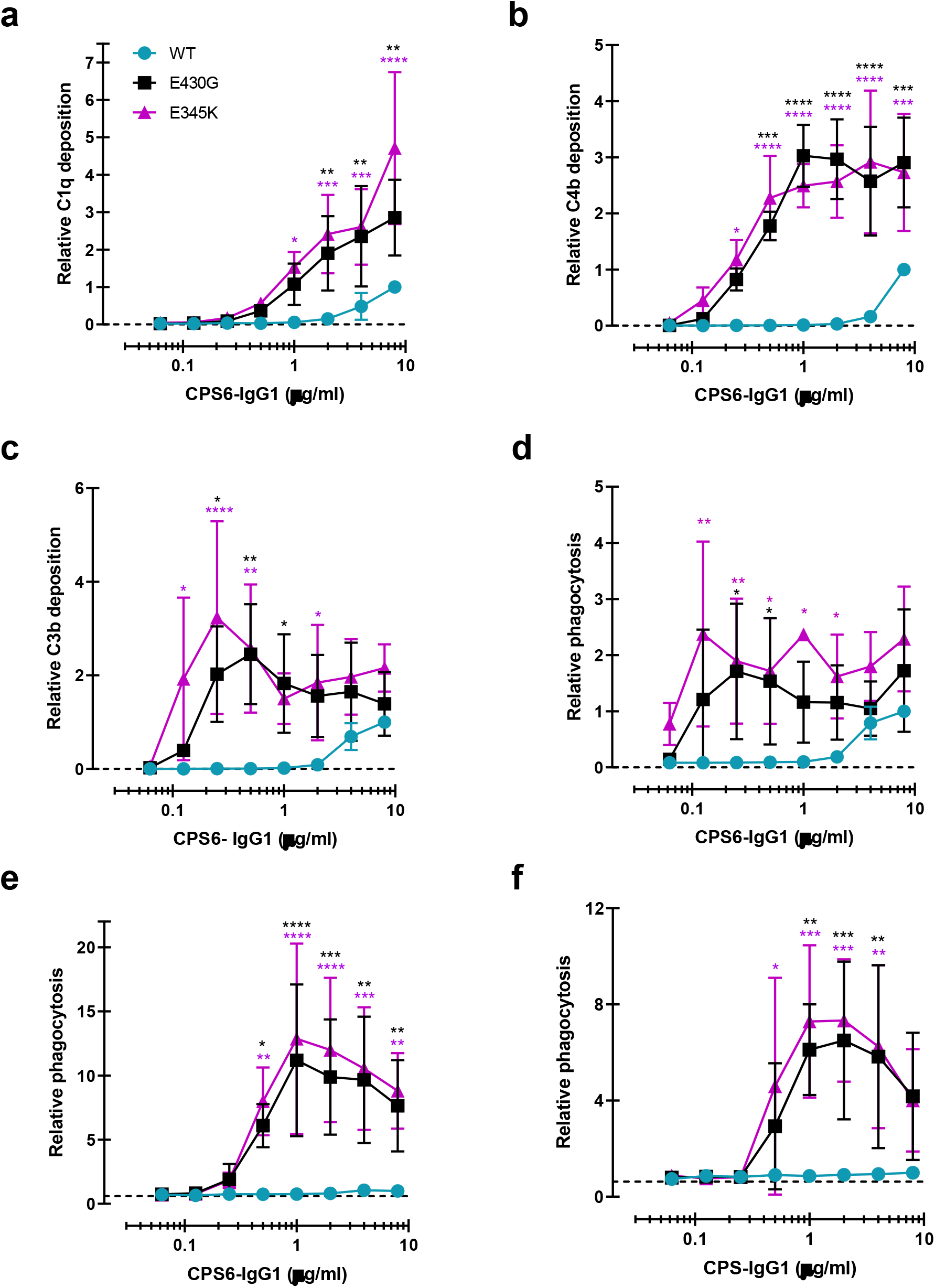
Enhanced Fc-Fc interactions strongly improves complement mediated phagocytosis of CPS6-IgG1 targeted *S. pneumoniae* serotypes. (a-c) Complement deposition on *S. pneumoniae* serotype 19A detected by flow cytometry after incubation with 2.5% IgG/IgM-depleted serum supplemented with CPS6-IgG1 (WT versus E430G and E345K variants). Detection of complement C1q (a), C4b (b) and C3b (c) deposition was done using a monoclonal anti-human C1q, C4d or C3d antibody, respectively. (d-f) Phagocytosis of fluorescently labelled *S. pneumoniae* serotype 19A (d), serotype 6A (e) and serotype 6C (f) by human neutrophils in the presence of 2.5% IgG/IgM-depleted serum supplemented with CPS6-IgG1-WT versus E340G and E345K variants. Bacterial uptake was quantified by flow cytometry as the mean fluorescence of the neutrophils. All data represent relative mean ± SD of three independent experiments and displayed by the relative fluorescence index compared to CPS6-IgG1-WT at 8 µg/ml. Dashed line represents background (no IgG) level. Two-way ANOVA was used to compare across dose-response curves at the various concentrations the differences between the WT and the E430G or E345K variants. When significant it was displayed as *P < 0.05; ***P < 0.001; ****P < 0.0001.

### CPS6-IgG1-E430G and CPS6-IgG1-E345K induce killing of *S. pneumoniae* by human neutrophils in normal serum

Having established that hexamer-enhancing variants of CPS6-IgG1 strongly induce phagocytosis of serogroup 6 and serotype 19A strains, we studied whether these antibodies also trigger intracellular killing of bacteria by neutrophils. To mimic the natural situation more closely, we now performed experiments in normal human serum (NHS) that contains pre-existing antibodies (62, 63) instead of using IgG/IgM-depleted serum as a complement source. This is important because 6B and 19A serotypes circulate in the healthy population and naturally occurring antibodies could thus play an additional role in phagocytosis by the mAbs. Previous experiments were repeated in the presence of normal serum as complement source. Indeed, E430G and E345K mutants also exhibited enhanced complement deposition (**Figure S7a,b**) and improved capacity to induce phagocytosis of serotype 6B (**Figure 5a**) and 19A (**Figure 5b**). It is noteworthy that the WT antibody performed better capacity to phagocytose bacteria in the presence of normal serum than in presence of IgG/IgM-depleted serum, potentially suggesting a collaboration of naturally occurring antibodies in pneumococcal internalization. In both cases the presence of active complement was required, as heat inactivation at 56°C completely abolished pneumococcal phagocytosis (**Figure 5a, b**). Next, to study whether antibody hexamerization facilitates bacterial killing following phagocytosis, bacteria were opsonized with CPS6-IgG1-WT or CPS6-IgG1-E430G/E345K variants in the presence of normal human serum before measuring the killing capacity of human neutrophils after 45 min by counting surviving colonies. Survival of serotypes 6B was strongly decreased in the presence of CPS6-IgG1-E430G or E345K in a dose-dependent manner, even achieving complete bacterial clearance at 3 μg/ml with E345K (**Figure 5c and S8a**), confirming that the killing of *S. pneumoniae* by neutrophils after uptake is an efficient process. When the effectiveness of the CPS6-IgG1 hexamer-enhancing antibodies to improve killing of its cross-reactive serotype 19A was assessed few, if any, bacteria were recovered (**Figure 5d and S8b**). These data show the functional positive consequences of antibody hexamerization on pneumococcal killing and their broader use among cross-reactive serotypes.

**Figure 5.**
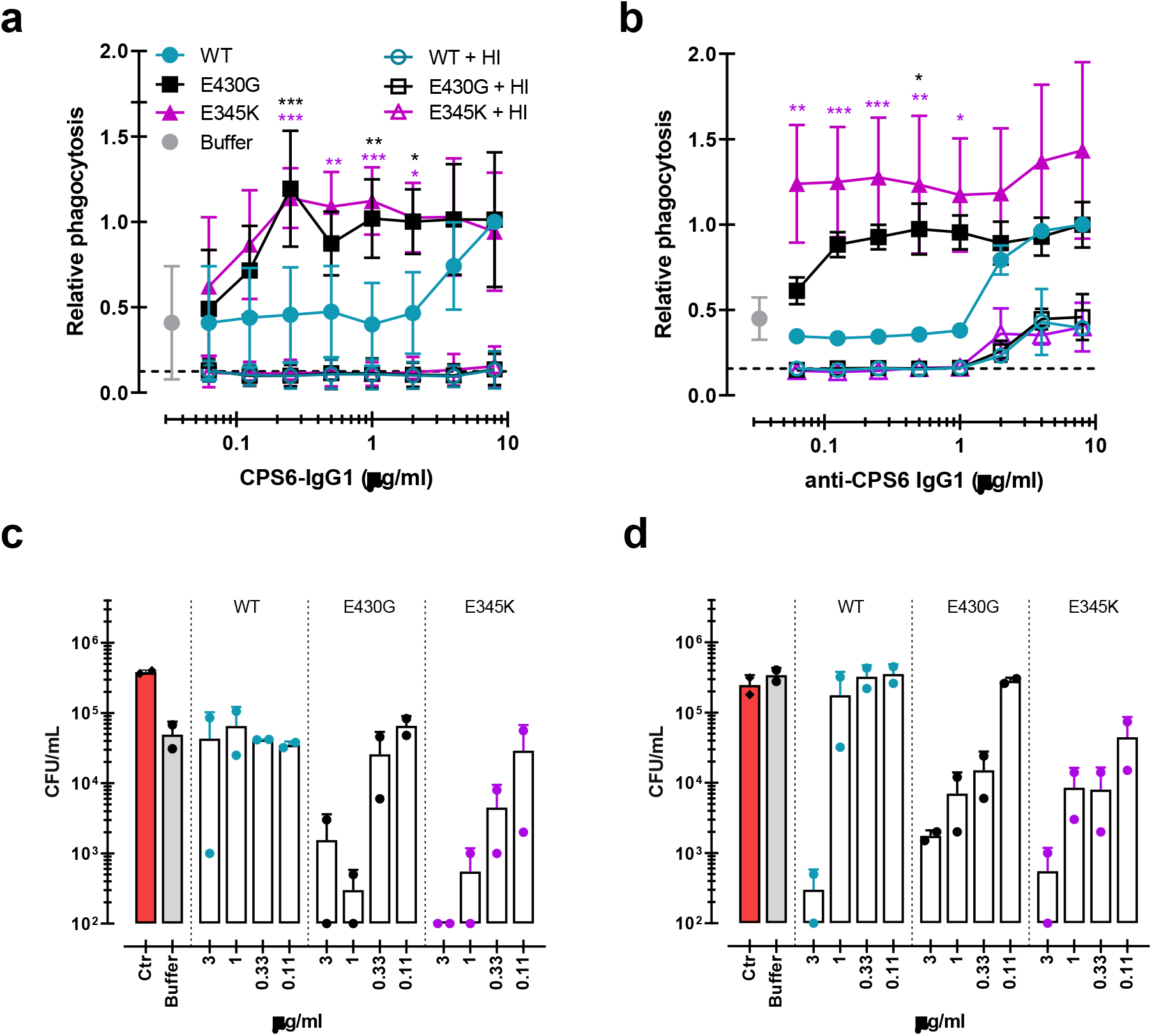
Monoclonal antibodies against *S. pneumoniae* capsule polysaccharide 6 can be modified for enhanced phagocytosis and killing by human neutrophils. (a-b) Phagocytosis by human neutrophils of fluorescently labelled *S. pneumoniae* serotype 6B (a) or serotype 19A (b) in the presence of 5% normal human serum (NHS) as complement source supplemented with CPS6-IgG1-wild-type (WT) versus E430G and E345K hexamerization-enhanced variants. Same conditions for phagocytosis but in the presence of 5% NHS without complement activity (heat inactivated, HI) is represented by empty coloured boxes. Bacterial uptake was quantified by flow cytometry as the fluorescence of the neutrophils. Data represent relative fluorescence mean index ± SD of three independent experiments compared to the highest CPS-IgG1-WT concentration tested (8 µg/ml). Dashed line represents background (no IgG) level and Buffer refers to the same condition with HPS (no IgG present). Two-way ANOVA was used to compare across dose-response curves at the various concentrations the differences between the WT and the E430G or E345K variants. When significant it was displayed as *P < 0.05; ***P < 0.001; ****P < 0.0001.(c, d) Phagocytic killing of *S. pneumoniae* serotype 6B (c) or serotype 19A (d) in the presence of 5% NHS and CPS6-IgG1-WT versus CPS6-IgG1-E430G or CPS6-IgG1-E345K mutant. Bacterial killing was determined after 45 minutes incubation with human neutrophils by counting colony formation units (CFU) on blood agar plates. Red bars (Ctr) represent initial bacterial inoculum, whereas grey bars (Buffer) represent bacterial killing when antibodies were omitted. Data represent the mean ± SD of two independent experiments with duplicate counting.

Overall, these results clearly show that mAb modification to induce hexamer formation on the bacterial surface potently increases complement-mediated *S. pneumoniae* phagocytosis and an effective intracellular killing.

### CPS6-IgG1-E345K engineered mAb for enhanced hexamerization protect mice against invasive pneumococcal infection

Following colonization of the nasopharynx, *S. pneumoniae* has the potential to invade the body and cause a broad spectrum of life-threatening diseases such as bacteremic pneumonia and meningitis. Transmission, colonization and invasion depend on the remarkable ability of this bacteria to evade the host inflammatory and immune responses (64). Hence, we investigated whether passive immunization with anti-capsular mAbs could protect mice against *S. pneumoniae* in a bacteremic pneumonia model. To this end, female BALB/c mice received an intraperitoneal injection of CPS6-IgG1-WT, CPS6-IgG1-E345K or PBS (**Figure 6a**). Three hours later, pneumonia was induced by intranasal challenge with pneumococcal serotype 6A (selected due to its higher virulence in mice (65)). Survival and bacterial loads in the blood were monitored for 7 days (**Figure 6 and S9**). The presence of mAbs in the bloodstream was confirmed by ELISA (**Figure S9c**). In PBS-treated controls, all mice developed bacteremia within 24 hours as evidenced by the presence of high loads of pneumococci in the bloodstream (**Figure 6b and S9A**). Passive administration with CPS6-IgG1-E345K effectively prevented bacterial dissemination to the bloodstream within the first 24h (**Figure 6b and S9A**). While 100 μg CPS6-IgG1-E345K could protect 60% of infected mice (12/20) from developing bacteremia, the same concentration of CPS6-IgG1-WT only reduced bacteremia in 27% of mice (4/15 mice) (**Figure 6b)**. A 3-fold higher concentration of CPS6-IgG1-WT was needed to achieve the same protection as CPS6-IgG1-E345K. Upon following survival, we observed that 95% (19/20) of PBS-treated mice succumbed due to the infection (**Figure 6c, S9a**). In contrast, 100 μg CPS6-IgG1-E345K could rescue 50% of mice (10/20). Again, we observed that the CPS6-IgG1-E345K mAb was more potent in protecting mice than CPS6-IgG1-WT. Additionally, mice weight regain was very much associated with the protective capacity of each mAb (**Figure S9b**).

**Figure 6.**
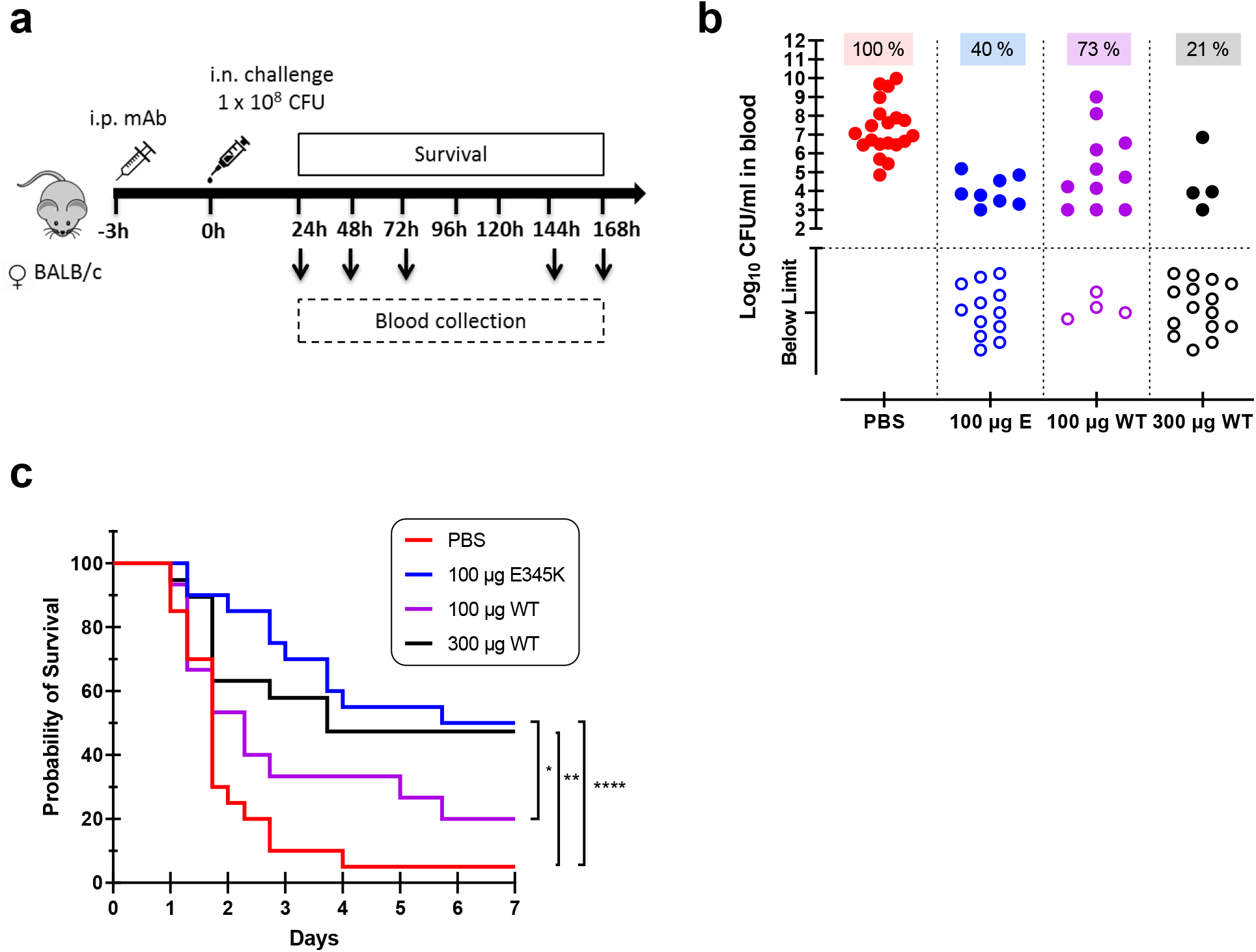
CPS6-IgG1-E345K mAb provides enhanced protection against invasive pneumococcal infection in mice. (a) Schematic representation of the infection model. Female BALB/c mice were passively immunized via intraperitoneal injection (i.p.) with PBS, 100 µg CPS6-IgG1-E345K, or 100 or 300 µg CPS6-IgG1-WT monoclonal antibody (mAb) 3 hours before infection. Mice were challenged intranasally (i.n.) with 1×10^8^ CFU *S. pneumoniae* serotype 6A in 50 µl PBS. Every day after challenge, blood was taken from the tail vein, serially diluted, and plated on blood agar plates for bacterial colony counting (CFU). (b) Bacteremia in mice blood 24h after bacterial challenge representing mAb capacity to control bacterial spread from lungs to the systemic circulation (PBS 100%, 20/20; 100µg E345K 40%, 8/20; 100µg WT 73%, 11/15; 300µg WT 21%, 4/15). Each symbol represents an individual mouse, closed symbols represent mice that developed bacteremia and open symbols represent mice below the threshold of CFU detection marked by the dotted line. Mice survival was monitored in parallel for 7 days (c). The data are combined from three separate experiments with five to eight mice for each treated group in each experiment resulting in 20 mice per group (only 15 mice for the 100 µg CPS6-IgG1-WT group). Statistical analysis was performed using Log-rank (Mantel-Cox) test and it was displayed when significant as *P ≤ 0.1, **P ≤ 0.01, or ****P ≤ 0.0001.

When repeating the same experiments in male BALB/c mice, we observed that 100 μg CPS6-IgG1-E345K and 300 μg CPS6-IgG1-E345K could not confer significant protection against bacteremia (**Figure S10**). However, these mAbs were able to control bacterial spread from lungs to the bloodstream within 24h after infection in the same proportion as in female mice (**Figure S10b**. Although this needs further investigation, these data potentially indicate gender differences in immune protection to pneumococcal infections.

Altogether, these data suggest that passive immunization with complement-enhancing monoclonal antibodies can prevent mice from developing bacteremia by encapsulated pneumococci.

### Promoting Fc-Fc interactions enhances complement-dependent phagocytosis via mAbs recognizing CPS3 and CPS8

Being aware of the fact that CPS6 mAbs can only target a small fraction of all circulating pneumococci, we also studied phagocytosis of mAbs against other capsular serotypes. We constructed IgG1 antibodies against capsule serotype 3 or 8, using variable region sequences of these mAbs available from literature (66, 67). Serotype 3 causes disease in both adults and children and has been associated with an increased risk of death compared to other pneumococcal serotypes (66, 68). Unlike serotype 3, serotype 8 is not included in PCV vaccines, and therefore its prevalence remains increasing (69, 70) world-wide. After verifying mAbs variants equal binding (**Figure S11a,b**), we studied complement mediated serotype 3 and 8 phagocytosis in the presence of 5% NHS and the CPS3 or CPS8 mAb variants. **Figure 7a,b**). Once more, we found that hexamerization-enhancing mutations highly enhanced phagocytosis of both serotypes 3 and 8 in a dose-dependent manner.

**Figure 7.**
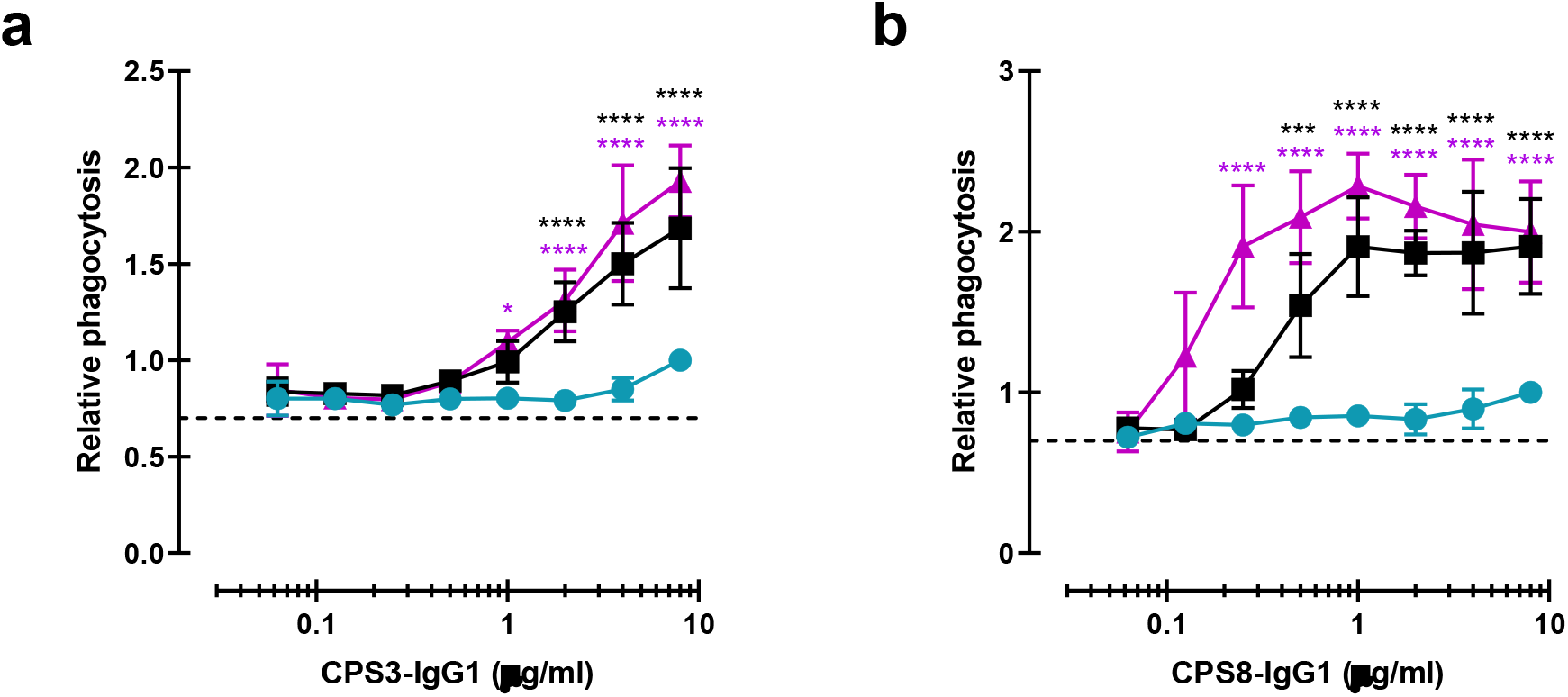
Monoclonal antibodies against CPS can be modified for enhanced complement mediated pneumococcal phagocytosis. (a, b) Phagocytosis by human neutrophils of fluorescence labelled *S. pneumoniae* serotype 3 (a) and serotype 8 (b) after incubation with 5% human normal sera supplemented with CPS-IgG1-WT, CPS-IgG1-E430G or CPS-IgG1-E345K variant. All data represent relative fluorescence mean index ± SD of three independent experiments compared to the highest CPS-IgG1-WT concentration tested (8 µg/ml). Dashed line represents background (no IgG) level. Two-way ANOVA was used to compare across dose-response curves at the various concentrations the differences between the WT and the E430G or E345K variants. When significant it was displayed as *P < 0.05; ***P < 0.001; ****P < 0.0001.

Overall, these results confirmed that antibodies directed against CPS could be used as therapy target as they can be modified to enhance hexamer formations, having strong positive influence on complement deposition and phagocytosis of highly virulent *S. pneumoniae* strains.

## Discussion

Here we studied whether monoclonal antibodies directed against the capsule of *S. pneumoniae* have the potency to induce bacterial killing via the human immune system. Although our data indicate that WT anti-capsular IgGs used in this study had a poor capacity to induce elimination of bacteria, the introduction of hexamer-enhancing mutations enabled such mAbs to induce potent immune clearance of *S. pneumoniae* infection, both *in vitro* and *in vivo*.

Immune-mediated killing of Gram-positive bacteria strongly depends on the capacity of human neutrophils to engulf bacteria via phagocytosis and kill them intracellularly (28, 71). Antibodies can enhance this process, by binding to the bacterial cell surface and stimulating FcγR uptake and/or activate the complement cascade to deposit C3-derived opsonins. Although the pneumococcal polysaccharide capsule is known to evade complement activation (by shielding epitopes that are hidden underneath the capsule (29)), our results support the idea that antibodies directed against the capsule can overcome capsular immune evasion and deposit complement opsonins on the encapsulated bacterium to stimulate phagocytosis. For the anti-capsular antibodies included in this study, we showed that monoclonal IgGs had a poor capacity to activate complement when expressed with wild-type Fc domains. This in sharp contrast to monoclonal IgGs recognizing wall teichoic acid (WTA) on the surface of *S. aureus* (72). Like the pneumococcal capsule, WTA is an abundant and exposed glycopolymer on the cell wall of *S. aureus* (73). For mAbs targeting WTA, we recently observed that WT IgG1 and IgG3 mAbs could induce complement activation and downstream phagocytosis in the absence of Fc-Fc enhancing mutations (72). At present it is unclear why these WT anti-capsular antibodies showed a poor capacity to induce complement activation on *S. pneumoniae*. The finding that these antibodies did not effectively recruit C1q, suggests they have a poor ability to establish Fc-Fc contacts needed for hexamer formation. Potentially, the capsule epitope is not an optimal antigen for IgG hexamer formation. For instance, molecular characteristics like antigen density, size and fluidity could affect hexamer formation (30). Also, the affinity of one Fab arm for the epitope could play a role since antibodies should bind with one Fab arm to enable hexamer formation (30). Whether the observed low capacity of wild-type CPS-specific mAb to induce IgG clustering can be extended to other anti-CPS or anti-pneumococcal antibodies would need further investigation. Potentially, our data may help to understand why certain capsular serotypes are poorly immunogenic. CPS antigen is poorly immunogenic in many individuals who are at the highest risk for the development of invasive pneumococcal disease and the use of protein conjugated vaccine has not been found in these patients to be more immunogenic than the purified polysaccharide-based vaccine (74). Since complement activation and deposition of C3-derived opsonins is important in driving adaptive immune responses to bacteria, for instance by enhancing B cell activation and antigen presentation (via complement receptors on B cells, APC’s and follicular dendritic cells) (75), a poor capacity to induce antibody clustering might also have a negative effect on vaccine efficacy.

Our findings also indicate that phagocytic uptake of *S. pneumoniae* via CPS-specific antibodies strongly depends on the presence of complement-induced opsonization, since we observed little to no uptake of antibody-labeled bacteria in the absence of complement. This reinforces the vital role of complement in pneumococcal clearance which is also demonstrated by the fact that patients with complement deficiencies are at high risk of *S. pneumoniae* infections (20, 76). Our finding that hexamer-enhancing mutations can boost complement-dependent killing of bacteria by itself is not new, since these same mutations have been demonstrated to potentiate complement-dependent killing of mAbs targeting *Neisseria gonorrhoeae* (77). However, complement-dependent killing of Neisseria is mediated by formation of membrane attack complex pores, which cannot act on Gram-positive bacteria (78). Our study provides an important proof-of-concept that hexamerization-enhancing mutations can potentiate opsonophagocytic killing of Gram-positive bacteria. This assumption is also supported by our *in vivo* bacteremic pneumonia mice model of infection. The presence of CPS-IgG1 mAbs potently prevented/delayed pneumococcal spread from lungs to the systemic circulation, resulting in effective clearing and, therefore, conferring protection against *S. pneumoniae* infection in female mice. The obtained protection was strongly increased by the use of engineered mAb CPS6-IgG1-E345K for enhanced hexamerization. Although this was not the goal of our study, the *in vivo* experiments also revealed that the monoclonal antibodies provided less protection against pneumococcal bacteremia in male than female mice. Clinical manifestation of the infection and mortality rate was comparable between male and female for the administered infectious dose. Dutch legislation encourages the verification of both genders in animal experiments, both for animal welfare and impact on translation to human diseases. Whether the immune responses to pneumococci and response to antibody therapies are truly different between males and females is beyond the scope of this paper but an important consideration for future therapy developments.

Overall, the presented work provides a proof-of-concept not only for the capacity of hexamer-enhancing mutations to improve anti-capsule mAb meditated immune system activation, but also for the potential use of monoclonal antibodies against encapsulated bacteria like *S. pneumoniae* when the existing therapies fail. In our study we demonstrated that the activity of mAbs directed against serogroup 6, one of the most prevalent serogroups worldwide (79, 80), could be effectively improved to prevent systemic pneumococcal infection. Furthermore, the fact that CPS6 mAbs likely cross-react with the highly invasive 19A strain, for which of invasive infection rates increased following PCV7 use worldwide (81), suggests the potential broader use of monoclonal antibodies. Similarly, hexamer-enhancing mutations could be used to improve the potency of recently discovered mAbs against other serotypes like CPS3 (82). Because complement is essential in immune protection against *S. pneumoniae* (26–28), the capacity of antibodies to induce complement activation could be exploited for effective antibacterial therapies while simultaneously avoiding the complications of antibiotic resistance. We are aware that the large variety pneumococcal CPS serotypes difficult their use as mAbs therapy for sufficiently coverage pneumococcal disease. Therefore, the most clinically useful scenario for the use of anti-pneumococcal CPS mAbs is to treat after symptoms onset. A cocktail composed of multiple anti-capsule mAbs would be very expensive unless, for instance, rapid test to identify the causative serotype can be developed to enable monovalent therapy. Monoclonal antibodies that react with a highly conserved surface antigens that elicit a portent immune response like histidine triad protein D (PhtD) (71) can be a promising tool for broad treatment against numerous pneumococcal serotypes. From the presented results, we anticipate this work will stimulate new routes to optimize antibody therapies against *S. pneumoniae* and other encapsulated bacteria.

## Materials and Methods

### Bacterial strains and fluorescent labelling

*S. pneumoniae* clinical isolates used in this study included strains from serotype 3, 6A, 6B, 6C, 8, 19A and 19F (kindly provided by Dr. J. Yuste, Centro Nacional de Microbiología, CNM-ISCIII, Madrid). Pneumococcal isolates were cultured on blood agar plates at 37°C in 5% CO_2_ or in Todd-Hewitt broth medium supplemented with 0.5% yeast (THY) extract to an optical density at 550 nm (OD_550_) of 0.6 and stored at −80°C in 10% glycerol as single use aliquots for further experiments. For generation of fluorescently labelled bacteria, strains were grown in THY, washed with PBS and incubated with fluorescein isothiocyanate (FITC; Sigma) (0.5 mg/ml) for 60 min on ice in dark. Bacteria were washed four times with PBS, resuspended in RPMI-0.05% human serum albumin (HSA) and stored as aliquots in 10% glycerol at −80°C.

### Isolation of human serum

Human blood from healthy volunteers was collected in plastic vacuette tubes (Greiner) with informed consent in accordance with the Declaration of Helsinki. Approval from the Medical Ethics Committee of the University Medical Center Utrecht was obtained (METC protocol 07-125/C approved on March 1, 2010). Clotting was allowed for 15 min and blood was centrifuge at 3,166 × g for 10 min at 4°C to collect serum. Sera of 20 donors was pooled and stored as single-use aliquots at −80°C as a source of complement and serum components. As an alternative complement source, the same NHS was depleted for IgG and IgM using a HiTrap Protein-G and Poros anti-IgM column in tandem on an Akta FPLC system (GE-Healthcare) as previously described (36).

### Human monoclonal antibody production

Human monoclonal antibodies were produced recombinantly in human Expi293F cells (Life Technologies) as described before (83), with minor modifications. Briefly, gBlocks (Integrated DNA technologies, IDT), containing codon-optimized variable heavy and light chain (VH and VL) sequences with an upstream KOZAK and HAVT20 signal peptide, were cloned into homemade pcDNA34 vectors, upstream the IgG heavy and kappa light chain constant regions, respectively, using Gibson assembly (New England Biolabs). The VH and VL sequences of the antibodies were derived from previously reported antibodies against CPS6 (84), CPS3 (66) and CPS8 (67), with some modifications (Table S1). Transfection of EXPI293F cells was performed using PEI (Polyethylenimine HCl MAX; Polysciences). After 4 to 6 days of transfection, IgG1 and IgG2 antibodies were isolated from cell supernatants using a HiTrap Protein A High Performance column (GE Healthcare), whereas IgG3 antibodies were isolated with a HiTap Protein G High Performance column (GE Healthcare). Antibodies were dialyzed in PBS, overnight at 4 °C, and filter-sterilized through 0.22 µm Spin-X filters. Antibodies were analyzed by size exclusion chromatography (GE Healthcare) and monomeric fractions were isolated in case of aggregation levels >5%. The concentration of the antibodies was determined by measurement of the absorbance at 280 nm and antibodies were stored at −20 °C until use.

Monoclonal IgG1 antibodies against CD52 (Alemtuzumab (85)), recombinantly expressed as WT and hexamer-forming RGY mutant (30), were obtained from Genmab (Utrecht, the Netherlands).

### Antibody binding and deposition of complement components on bacterial surface

Antibody binding and complement C1q, C4b and C3b deposition was assessed on FITC-labelled strains in RPMI-HSA using flow cytometry as previously described (86). For anti-CPS binding, bacteria were incubated with antibody for 20 min at 4°C, washed and incubated for another 30 min at 4°C with APC-labelled donkey-anti-human-IgG (Jackson ImmunoResearch Europe Ltd.). For complement deposition assays, 2.5% IgG/IgM-depleted serum or 5% NHS was used. Bacteria were incubated with a concentration range of 2-fold serial diluted mAb against CPS (starting at 10 μg/ml) plus a fixed serum concentration for 30 min at 37°C. Subsequently, bacteria were washed with buffer and incubated with specific monoclonal antibodies for human C1q, C4b or C3b (all at 1 μg/ml; Quidel) for 30 min at 4°C. Complement components binding was detected with Alexa-conjugated F(ab’)2-goat anti-mouse IgG (H + L) (2 μg/ml; Jackson ImmunoResearch Europe Ltd) after 30 min incubation at 4°C. Samples were washed, fixed with 1% ice-cold paraformaldehyde (PFA) and measured in a flow cytometer (BD FASCVerse). Data were analyzed by FlowJo software and results are presented as relative mean fluorescence index (MFI), compared to the highest concentration of the CPS-IgG-WT.

### Opsonophagocytosis and killing of *S. pneumoniae* by neutrophils

Human neutrophils were freshly isolated from healthy donor blood using the Ficoll-Histopaque gradient method already described (44, 87). Phagocytosis assay was performed in a round-bottom 96-well plate and neutrophil-associated fluorescent bacteria were analysed by flow cytometry. FITC-labelled *S. pneumoniae* strains were opsonized by pre-incubated with 2-fold serial dilutions of mAb in 5% NHS, or 2.5% IgG/IgM-depleted serum as complement source, in RPMI-HSA or as control in 5% heat-inactivated NHS (30 min at 56°C) for 20 minutes at 37°C. Subsequently, neutrophils were added in a 1:10 cell to bacteria ratio and phagocytosis was allowed for 30 min at 37°C on a shaker (650 rpm). Ice-cold 1% PFA in RPMI-HSA was added to stop the reaction. Samples were measured by flow cytometry, and mean fluorescence values determined for gated neutrophils (87). Results are presented as relative mean fluorescence index (MFI) compared to the highest concentration of CPS-IgG-WT. When neutrophil opsonophagocytic killing capacity was assessed, similar procedure was performed with some modifications based on previously described method (88). Bacteria were opsonized during 20 min in a round-bottom 96-well plate as described above. For each condition, the mixture was subsequently transferred to sterile none-siliconized 2 ml tubes (Sigma) with neutrophils (1 × 10^7^ cells) in a 1:1 ratio in 100 μl volume and the phagocytosis process was prolonged to 45 min at 37°C on a shaker (650 rpm) to ensure intracellular bacterial killing. To release the internalized bacteria, the neutrophils were lysed for 5 min with ice cold 0.3% (wt/vol) saponin (Sigma-Aldrich) in sterile water. Samples were then serially diluted in PBS and plated onto blood agar plates in duplicate. CFUs were counted after overnight incubation at 37°C 5% (vol/vol) CO2 incubator, and percentage survival was calculated and compared with inoculum.

### Fc-III peptide

The inhibitory Fc-binding peptide (DCAWHLGELVWCT)(46) and a scrambled version of Fc-binding peptide sequence, Scr (WCDLEGVTWHACL), were both synthesized by Pepscan (Lelystad, The Netherlands). A fixed concentration (10 μg/ml) was added together with bacteria and sera, before the addition of human neutrophils.

### Native mass spectrometry

Native MS experiments were performed a standard Exactive Plus EMR Orbitrap instrument (ThermoFisher Scientific). Before analysis of protein samples, buffers were exchanged to 150 mM ammonium acetate (pH 7.5) through six dilution and concentration steps at 4 °C using Amicon Ultra centrifugal filters with 10 kDa molecular weight cutoff (Merck). For experiments studying the IgG-binding properties of Fc-III and Scr, 1 µM of anti-CD52 IgG1 was incubated with 4 µM Fc-III or 10 µM Scr for at least 15 min at RT. Anti-CD52 IgG1-RGY hexamers were measured at a total IgG1 concentration of 2 µM in the presence or absence of 40 µM Fc-III or Scr. The incubation step with the peptides was proceeded for at least 90 min at 37 °C due to the relatively slow disassembly rate of solution-formed IgG1-RGY hexamers. Samples were loaded into gold-coated borosilicate capillaries (prepared in-house) for direct infusion from a static nano-ESI source. Deconvoluted mass spectra were generated by Bayesian deconvolution using UniDec (89).

### Microscopy

Microscopy image of pneumococcal internalization by human neutrophils were performed after incubation of *S. pneumoniae* 6B in the presence of 5% IgG/IgM-depleted serum supplemented with CPS6-IgG1 E430G (8 µg/ml), as previously describe. Samples were prepared by cytospin (Shandon) and stained with Diff-Quick. Pictures were taken with a Sony Nex-5 camera mounted without lens on an Olympus BX50 microscope using a100X/1.25 oil objective to visualize cytoplasmic internalization.

### Mice

BALB/c mice from Envigo (Horst, The Netherlands), 8-12 weeks old, that matched for age and sex in each experiment were used. The animals were maintained on a 12-hour light/dark cycle in a room maintained at a mean temperature of 21 ± 2°C with a relative humidity of 50 ± 20%. Drinking water and food pellets were provided *ad libitum*. Animal experiments were performed at the infection Unit of the Central Animal Facility at Utrecht University and handled according to the institutional and national guidelines for the use and care of laboratory animals. The study was approved by the institutional Animal Care and Use Committee (AVD1150020172204).

### Pneumococcal infection in mice

The pneumococcal infection model has been described before (84, 90). In brief, mice were passively immunized intraperitonealy (i.p.) with 200 μl of antibody in PBS, 3h before challenge. Mice were anesthetized with inhaled isoflurane at 3% and challenged by intranasal (i.n.) route with 1 × 10^8^ CFU pneumococci serotype 6A in 50 μl PBS. Every twenty-four hours after challenge animals were weighted, scored by 2 independent researchers and blood was taken from tail vein. When mice exhibited severe signs of disease they were sacrificed according to the national guidelines. Blood was serially diluted, and plated on Colombia agar containing 5% sheep blood (Sanofi Diagnostics Pasteur, Marnes-la-coquette, France). Plates were incubated overnight at 37°C, and pneumococcal colonies were counted. Animals that survived for 7 day were considered protected.

### Enzyme-linked immunosorbent assays (ELISA) for IgG levels

Human antibody concentrations in mouse sera after passive immunization was measured by ELISA using 96-well half-area plates (Corning, #3690) coated with 2 µg/ml sheep anti-Human-IgG (ICN, Affi-pure) in 0.1M Carbonate buffer pH 9.6 for at least 18 h and blocked with 4% bovine serum albumin (BSA; Serva) in PBS+0.05% Tween-20. Captured IgG was detected with peroxidase-conjugated F(ab’)2-Goat-anti-Human-IgG-Fc antibody (Jackson) for 1 h at 37°C. Reaction was developed using a freshly prepared TMB substrate solution and stopped with 1M Sulfuric acid before determining the OD_450_ using a microtiter plate reader (iMark mic, Bio-Rad).

### Statistical analysis

Statistical analysis was performed with GraphPad Prism software (version 8.3). All data are presented as means ± SD from at least two independent experiments as indicated in the figure legends. Differences in the efficacy of the hexamerization-enhanced variants and the WT antibody across the different concentrations of the dose-response curves were assessed by two-way ANOVA using Dunnett’s multiple comparisons test, with individual variances computed for each comparison. For mice survival protection capacity between groups Log-rank (Mantel-Cox) test was used. When two groups were compared unpaired two-tailed t-test was performed.

## Acknowledgments

This work was supported by the Netherlands Organization for Scientific Research (NWO) through the European Union’s Horizon 2020 research programs H2020-MSCA-IF (#798032, to LA), the TTW-NACTAR Grant #16442 (to AJRH and SHMR), and ERC Starting grant (#639209, to SHMR). The authors thank Gestur Vidarsson (Sanquin, Amsterdam) and Paul Parren for scientific input and Alexandra Terry (Genmab) for the generated illustrations.

