## Supplemental figures for "Promoting Fc-Fc interactions between anti-capsular antibodies provides strong immune protection against *Streptococcus pneumoniae*"

### SUPPLEMENTARY INFORMATION - Promoting Fc-Fc interactions between anti-capsular antibodies provides strong complement-dependent immune protection against *Streptococcus pneumoniae*

Leire Aguinagalde[1], Maurits A. den Boer[2,3], Suzanne M. Castenmiller[1], Seline A. Zwarthoff[1], Carla J.C Gosselaar-de Haas[1], Piet C. Aerts[1], Frank J. Beurskens[4], Janine Schuurman[4], Albert J.R. Heck[2,3], Kok P.M. van Kessel[1] and Suzan H.M. Rooijakkers[1*]

[1]*Medical Microbiology, University Medical Center Utrecht, Utrecht University, The Netherlands*
[2]*Biomolecular Mass Spectrometry and Proteomics, Bijvoet Center for Biomolecular Research and Utrecht Institute for Pharmaceutical Sciences, Utrecht University, Padualaan 8, 3584 CH Utrecht, The Netherlands*
[3]*Netherlands Proteomics Center, Padualaan 8, 3584 CH Utrecht, The Netherlands*
[4]*Genmab, Utrecht, The Netherlands*

#### Figure S1

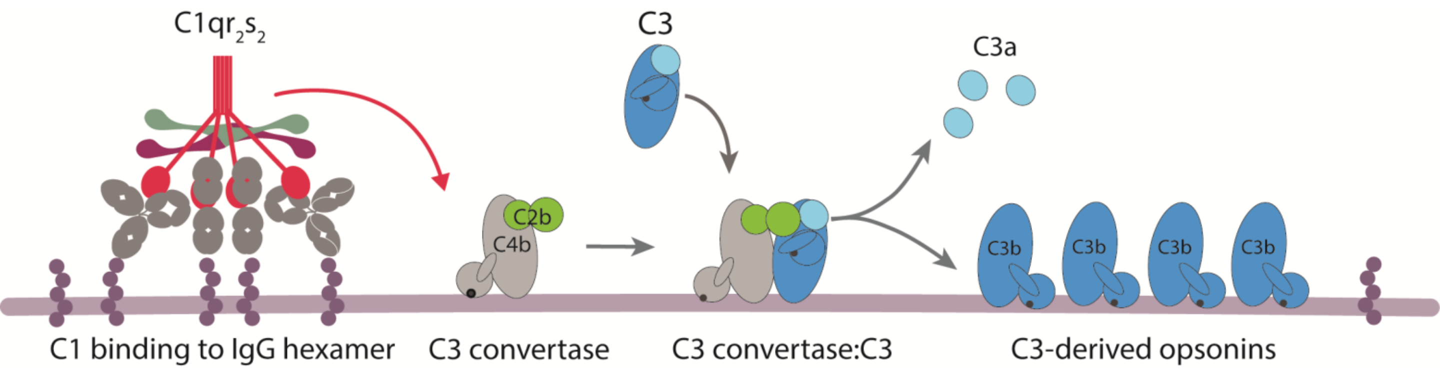


**Supplemental figure 1. Overview of complement classical pathway activation.** The binding of C1 complex ($C1qr_2s2$) to bacterium-bound antibodies triggers the activation of the classical pathway. C1 complex consists of an antibody recognition unit (C1q) and four associated proteases (2 copies of C1r and C1s). C1s will cleave C4 and C2 to generate a surface bound C3 convertase (C4b2b) that catalyses the rapid deposition of C3b molecules onto the target surface. C1 binding to antibodies occurs via the globular heads of C1q. Due to a low affinity of each gC1q domain for an IgG, avid binding of C1q requires clustering of surface-bound IgGs into an ordered hexamer that is held together via non-covalent Fc-Fc contact between neighbouring antibodies.

### Figure S2

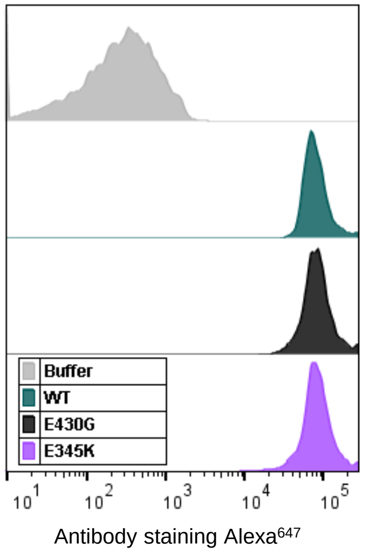


**Supplemental figure 2**. Representative flow cytometry histogram overlay showing equal binding of CPS6-IgG1 wild-type (WT), CPS6-IgG1-E430G and E345K mutants (4 µg/ml) to pneumococcal serotype 6B. Buffer control (grey) represents bacteria in absence of antibodies.

### Figure S3

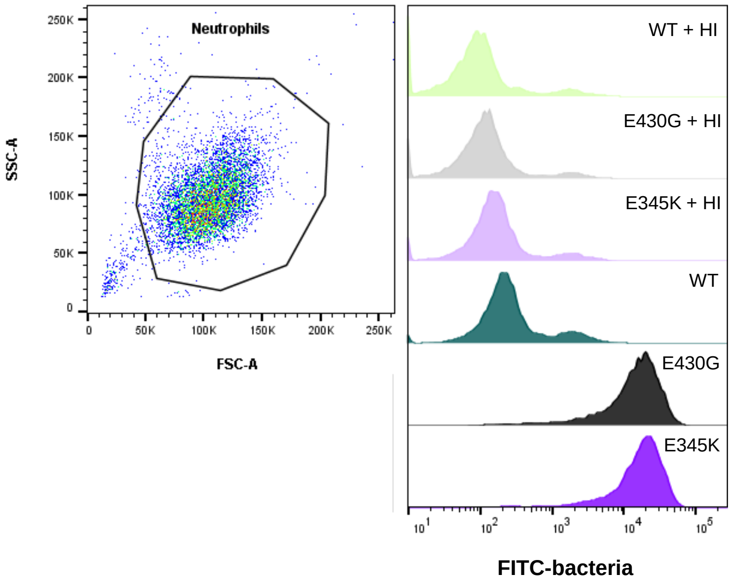


**Supplemental figure 3**. Representative flow cytometry analysis of pneumococcal serotype 6B phagocytosis by neutrophils. (*left*) Scatter plot of neutrophils and indicated gate setting for analysis. (*right*) Histogram overlay showing the FITC fluorescence of gated neutrophils in the presence of CPS6-IgG1-WT, CPS6-IgG1-E430G or E345K mutants (4 µg/ml) plus 2.5% heat-inactivated (HI; no active complement) or normal IgG/IgM-depleted serum (dNHS).

### Figure S4

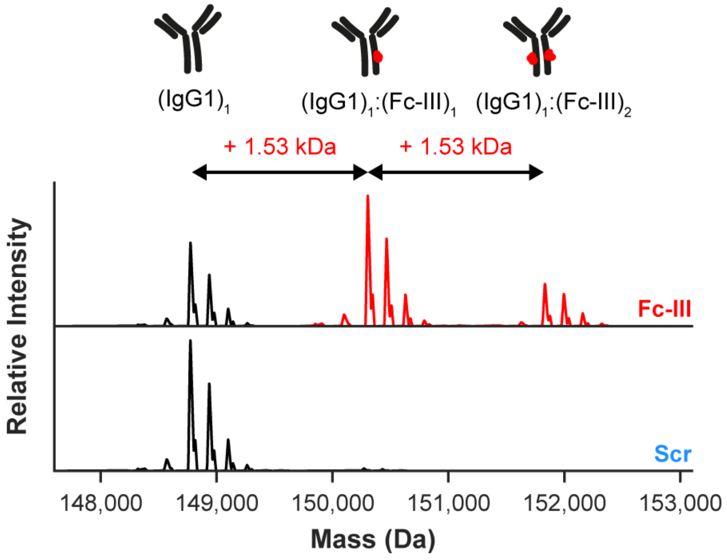


**Supplemental figure 4.** Deconvoluted native mass spectra showing that the masses of anti-CD52 IgG1 glycoforms (black) are shifted when incubated with Fc-III (red), but not with Scr (blue). This shift corresponds to binding of one or two copies of Fc-III to a single IgG1 molecule.

### Figure S5

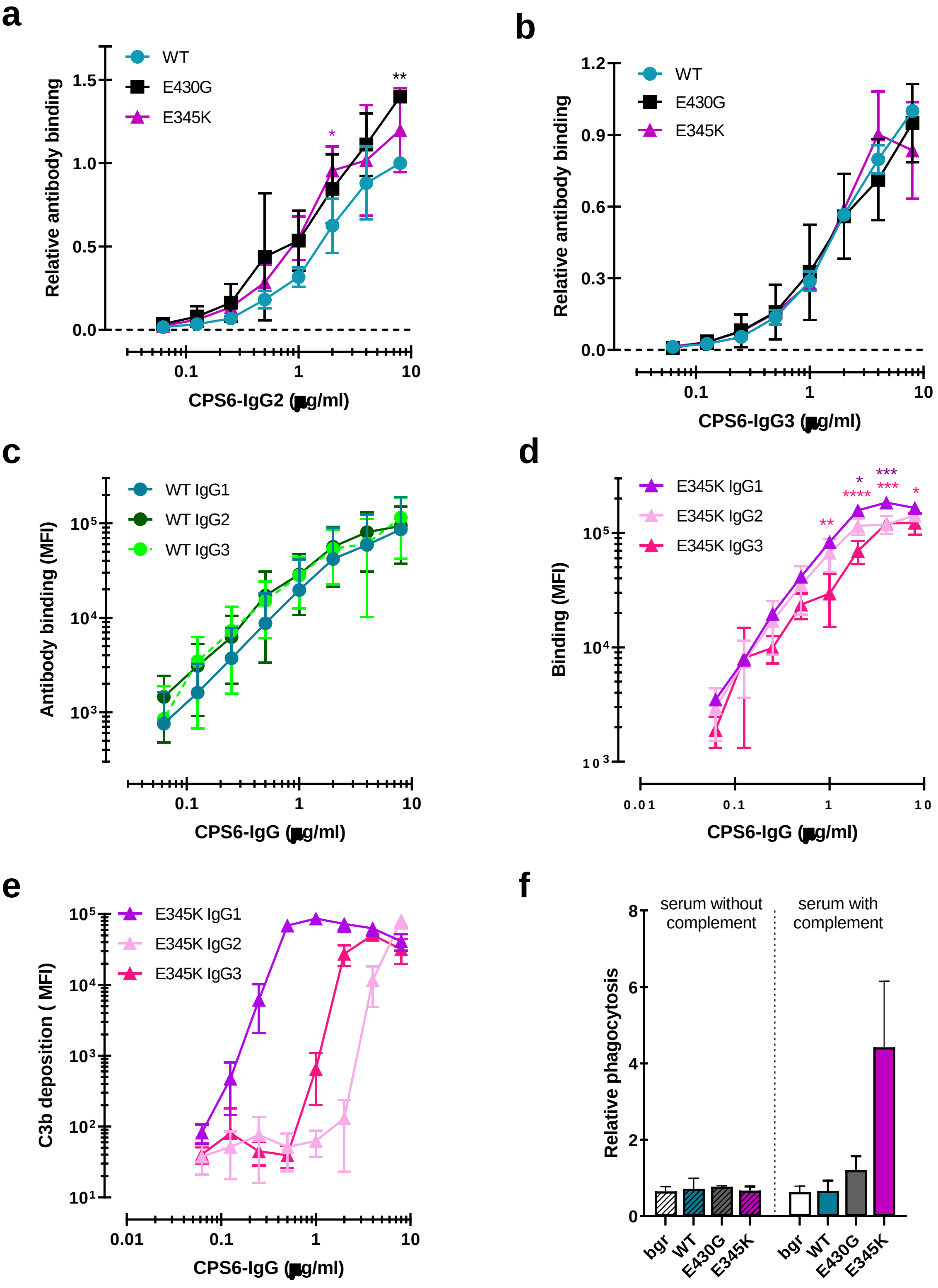

a
WT
E430G
E345K
Relative antibody binding
CPS6-IgG2 (μg/ml)
b
WT
E430G
E345K
Relative antibody binding
CPS6-IgG3 (μg/ml)
c
WT IgG1
WT IgG2
WT IgG3
Antibody binding (MFI)
CPS6-IgG (μg/ml)
d
E345K IgG1
E345K IgG2
E345K IgG3
Binding (MFI)
CPS6-IgG (μg/ml)
e
E345K IgG1
E345K IgG2
E345K IgG3
C3b deposition ( MFI)
CPS6-IgG (μg/ml)
f
serum without complement
serum with complement
Relative phagocytosis
bgr
WT
E430G
E345K
bgr
WT
E430G
E345K

**Supplemental figure 5**. Comparison of IgG2 and IgG3 anti-CPS6 antibodies. (a, b) Binding of CPS6-WT, E430G and E345K mutants as IgG2 (a) or IgG3 (b) variants to *S. pneumoniae* serotype 6B. Data are expressed relative to the 8 µg WT value and presented as means ± SD of three independent experiments. Dashed line represents background (no IgG) level. (c, d) Comparison of IgG1, IgG2, and IgG3 WT (c) and their E345K variants (d) binding to *S. pneumoniae* serotype 6B, expressed as mean ± SD of geometric mean fluorescence intensities (MFI) from three independent experiments. (e) Comparison of the complement C3b deposition of the IgG1, IgG2, and IgG3 E345K variants onto serotype 6B pneumococci. Data are mean ± SD of geometric mean fluorescence intensities (MFI) from three independent experiments(f) Phagocytosis of S. *pneumoniae* serotype 6B by CPS6-IgG3-E345K in 2.5% IgG/IgM-depleted serum without (HI) or with an active complement system presented relative to the MFI of IgG3-WT at 8 µg/ml. Bar graphs represent the relative phagocytosis values at 4 µg/ml of IgG3 WT, E430G and E345K mutant in 2.5% IgG/IgM-depleted serum without (striped bars) or with an active complement system (non-striped bars). Grey bars (Buffer) represent bacteria uptake by human neutrophils when antibodies were omitted. Data are mean ± SD of three independent experiments. Two-way ANOVA was used to compare across dose-response curves at the various concentrations the differences between the WT and the E430G or E345K variants. When significant it was displayed as *P < 0.05; ***P < 0.001; ****P < 0.0001.

### Figure S6

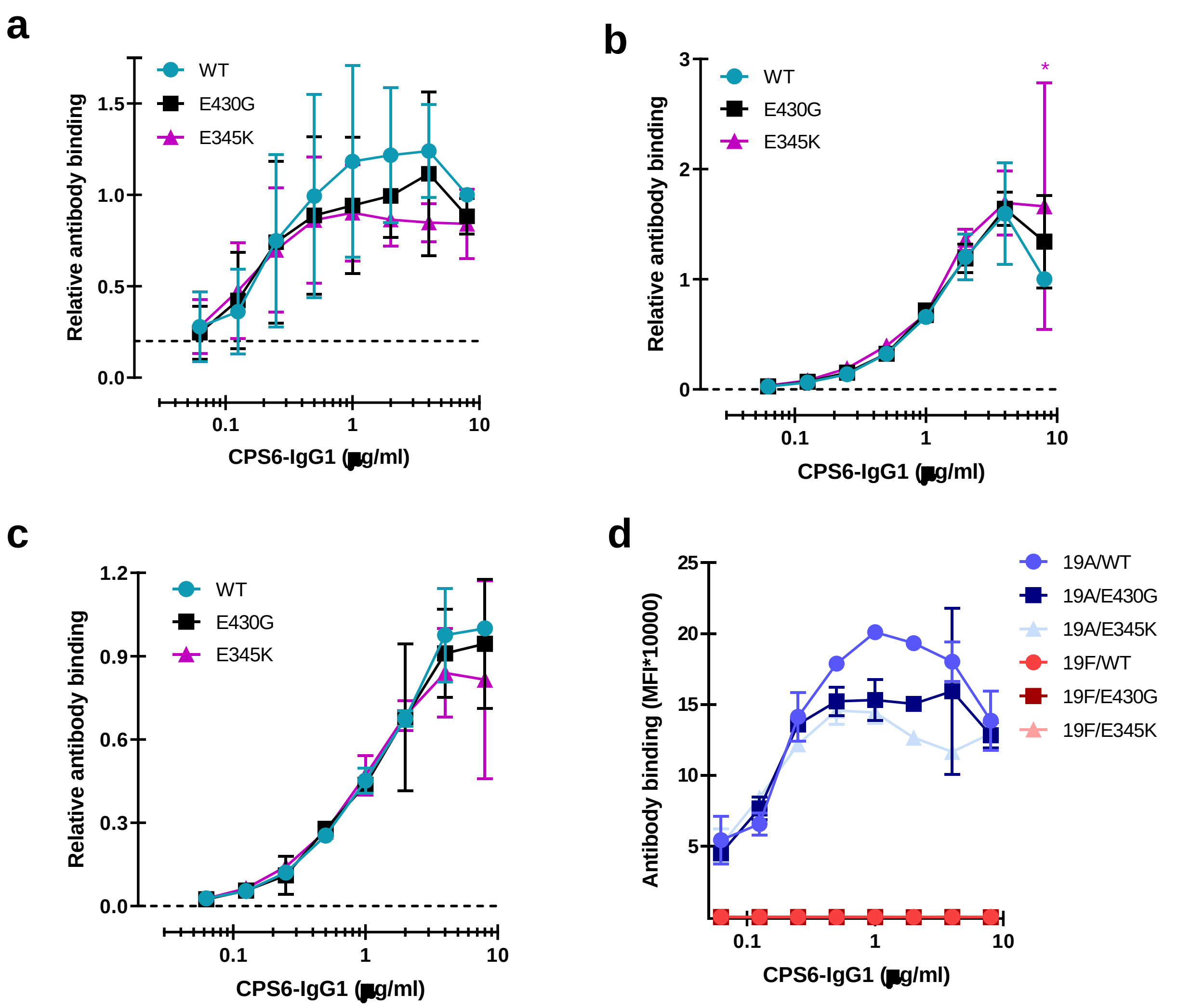


**Supplemental figure 6.** (a-d) Binding of CPS6-IgG1 WT or hexamer-enhancing mutations, E430G and E345K, to *S. pneumoniae* serotype 19A (a), 6A (b), 6C (c) but not to serotype 19F (d). (a-c) Data represent relative fluorescence mean index ± SD of three independent experiments compared to the highest CPS-IgG1-WT concentration tested (8 µg/ml). Two-way ANOVA was used to compare across dose-response curves at the various concentrations the differences between the WT and the E430G or E345K variants. When significant it was displayed as *P < 0.05; ***P < 0.001; ****P < 0.0001. (d) Data are expressed as mean ± SD of mean geometric fluorescence intensities (MFI) of two independent experiments. Two-way ANOVA was used to compare across dose-response curves at the various concentrations the differences between the WT and the E430G or E345K variants. When significant it was displayed as *P < 0.05; ***P < 0.001; ****P < 0.0001.

#### Figure S7

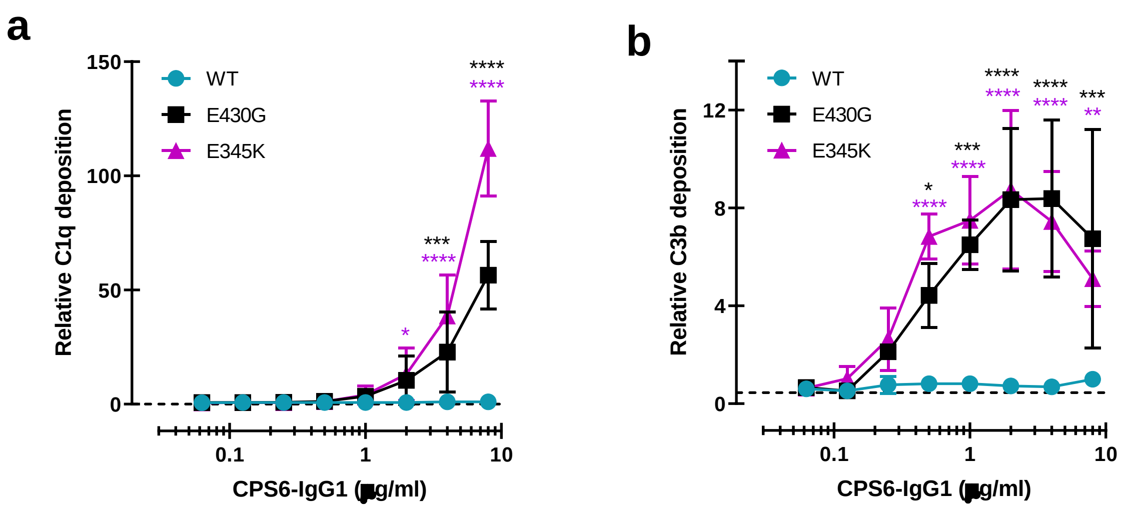


**Supplemental figure 7.** (a) C1q binding and (b) C3b deposition onto serotype 6B surface after incubation of bacteria with CPS6-IgG1 antibody variants, in presence of 5% human normal sera and detected with a monoclonal murine anti-human C1q and C3d antibody, respectively, by flow cytometry. Data represent mean ± SD of three (a) or two (b) independent experiments and are expressed relative to the highest CPS6-IgG1-WT concentration tested (8 μg/ml). Dashed line represents background (no IgG) level. Two-way ANOVA was used to compare across dose-response curves at the various concentrations the differences between the WT and the E430G or E345K variants. When significant it was displayed as *P < 0.05; ***P < 0.001; ****P < 0.0001.

### Figure S8

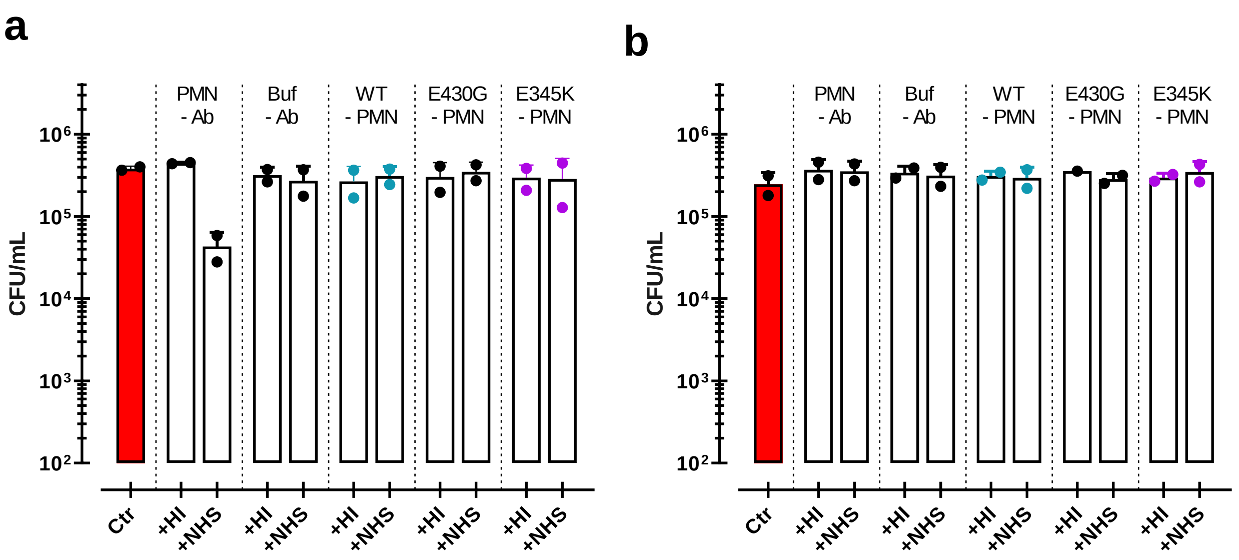


**Supplemental figure 8.** (a, b) Controls used for phagocytic killing of *S. pneumoniae* ST6B (a) or serotype 19A (b) in the presence of 5% heat inactivated (+HI) or normal human sera (+NHS). Controls in the absence (-Ab) of mAb contain only neutrophils (PMN) or only buffer (Buffer). Controls of only CPS6-IgG1-WT versus CPS6-IgG1-E430G or CPS6-IgG1-E345K mutant are without neutrophils (-PMN). Bacterial killing was determined after 45 minutes incubation with human neutrophils by colony formation unit (CFU) counting in blood agar plates. Red bars represent initial bacterial inoculum for the experiment.

### Figure S9

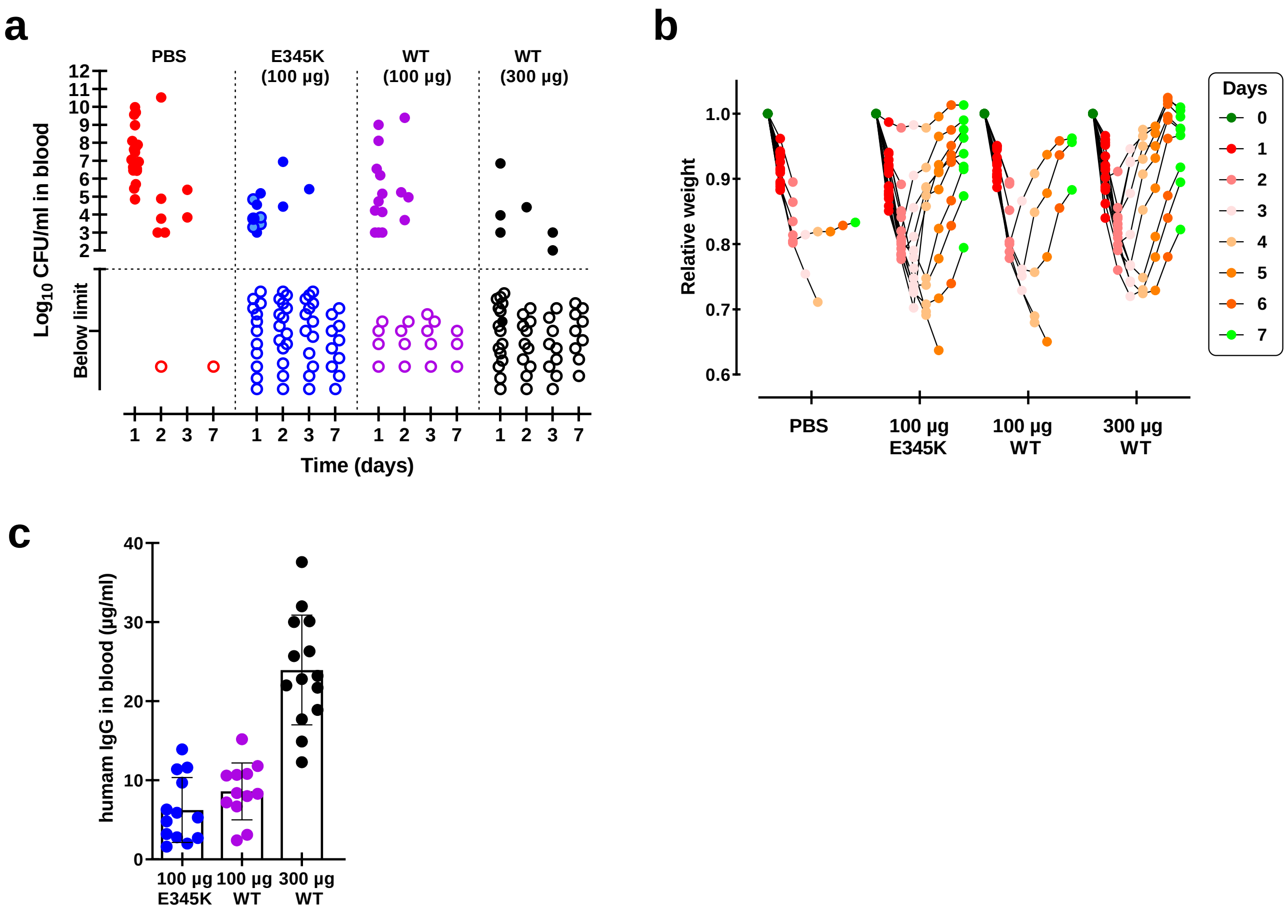


**Supplemental figure 9.** Female BALB/c mice infection model**.** (a) Bacterial counts in blood collected from surviving female mice on day 1, 2, 3 and 7 after challenge with 1x$10^8$ CFU *S. pneumoniae* 6A in order to compare protection capacity against bacteremia in mice passively immunized with only PBS, CPS6-IgG1 wild-type (WT) or with the E345K hexamerization-enhanced mutant. Each symbol represents an individual mouse, and the dotted line marks threshold of detection (2 log10 CFU/ml). (b). Weight of the individual mice as determined each day following the passive immunization for each treatment group. Data are expressed as relative weight values up to 7 days after bacteria challenge compared to the initial weight of each animal at the start of the experiment on day 0. (c) Human IgG detected by ELISA in mice sera 24h after passive-immunization with 100 or 300 µg CPS6-IgG1-wild-type (WT) or CPS6-IgG1-E345K expressed as mean µg/ml ± SD. All data are pooled from three independent experiments with finally 20 mice for PBS, 100 µg CPS6-IgG1-E345K, and 300 µg CPS6-IgG1-WT groups, and 15 mice for 100 µg CPS6-IgG1-WT group.

### Figure S10

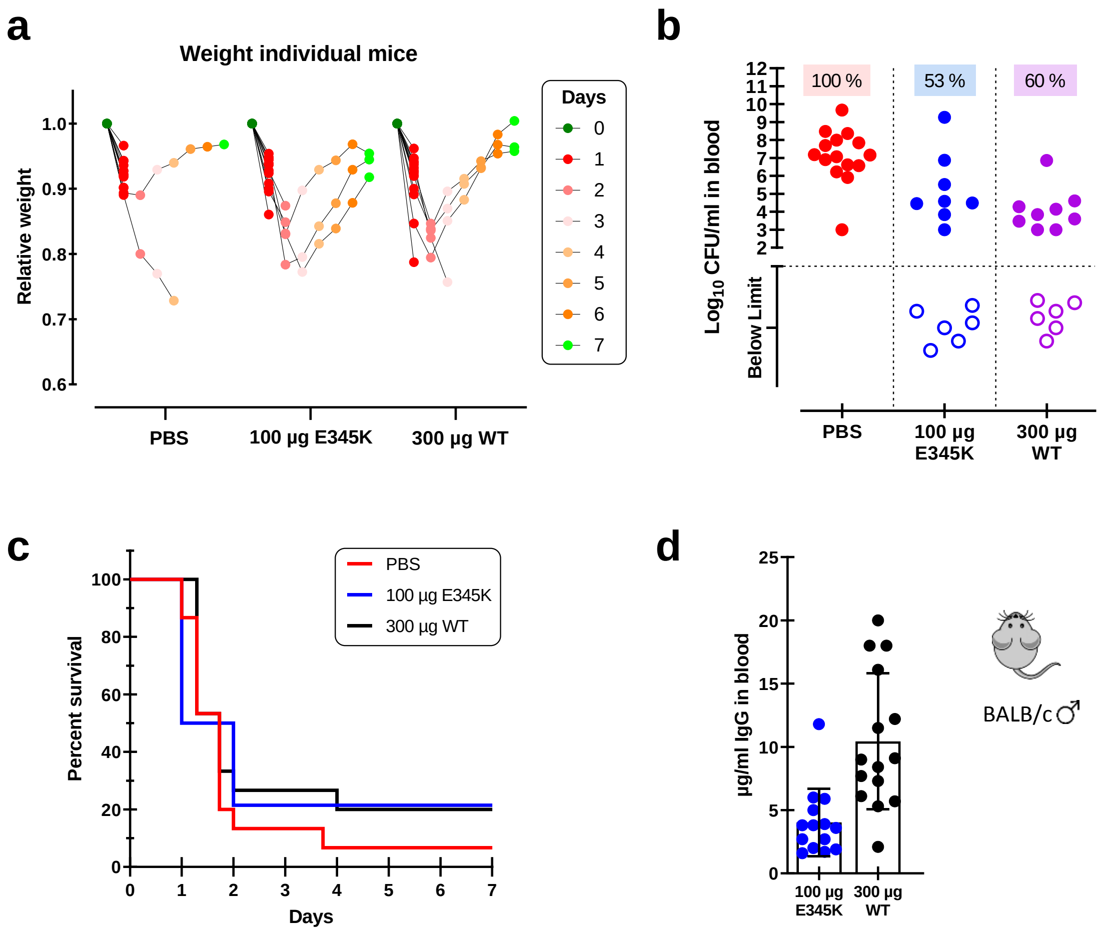


**Supplemental figure 10. Protective efficacy in male mice of engineered CPS6-IgG1-E345K.** (a) Male BALB/c mice (n=15 per group) passively immunized with PBS, 300 µg CPS6-IgG1-wild-type (WT) or 100 µg/ml CPS6-IgG1-E345K (E) were challenge with $1x10^8$ cfu *S. pneumoniae* serotype 6A and monitored for weight loss (a), bacteremia (b) and survival (c) for 7 days. (a) Data are expressed as relative weight values compared to the initial weight of each animal at the start of the experiment at day 0. (b) Dotted lines mark threshold of detection of bacteremia in mice blood 24h after bacteria challenge, representing mAb capacity to control bacterial spread from lungs to the systemic circulation. Each symbol represents an individual mouse, and closed dots represent mice that developed bacteremia. (c) Mice survival was monitored in parallel for 7 days. (d) Human IgG titers measured by ELISA in mice sera collected 24 hours after passive immunization with monoclonal antibody.

### Figure S11

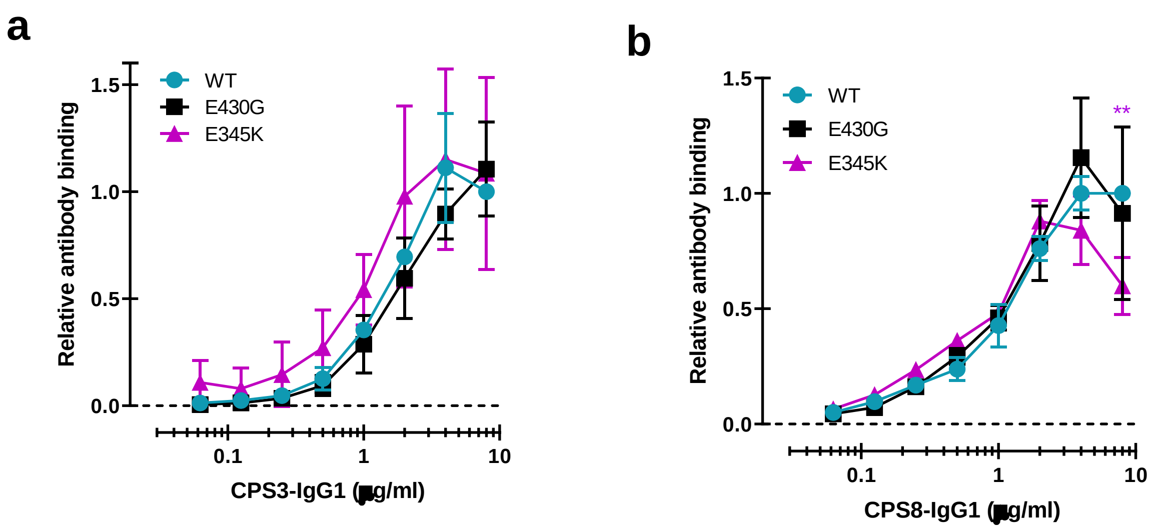


**Supplemental figure 11.** (a, b) Equal binding of CPS3-IgG1-WT (a) or CPS8-IgG1-WT (b) antibody, and their E430G and E345K variants to *S. pneumoniae* serotype 3 and 8, respectively. All data represent relative fluorescence mean index ± SD of three independent experiments compared to the highest CPS-IgG1-WT concentration tested (8 µg/ml). Dashed line represents background (no IgG) level. Two-way ANOVA was used to compare across dose-response curves at the various concentrations the differences between the WT and the E430G or E345K variants. When significant it was displayed as *P < 0.05; ***P < 0.001; ****P < 0.0001.

**Supplemental Table 1: Protein sequences used for antibody production.** Variable and constant heavy and light chain protein sequences used for antibody production. The residues E345 and E430 are highlighted in light grey and dark grey, respectively. The adapted amino acids for this study are highlighted in red.

| | **Sequence** |
|---|---|
| **Variable heavy chain** | |
| CPS6-IgG | EVQLVESGGGLVTFGGSLTLSCAASGFTFSRAWLTWVRQAPGGGLEWVGRILRMADGGATDYAATVKGRF<br>TISRDDSKNTVYLHMNNLKTEDTAVYYCANENFWRLDNWGQGTLVTVSS |
| CPS8-IgG(28H11) | EVKLLESGGGLVQPGGSLKLSCAASGFDFSRYWMSWVRQAPGKGLEWIGEINPDSSTINYTPSLKDKFIISR<br>DNAKNTLYLQMSKVRSEDTALYYCARPWFRYFDVWGAGTTVTVSS |
| CPS3-IgG(5F6) | QVQLQQPGAELVRPGASVKLSCKASGYTFTSYWINWMKQRPGQG<br>LEWIGVIDPSDSETHYNQMFKDKATLTVDKSSSTAYMQLSSLTSEDSAVYYCARRGDDGYTPAWFAYWGQ<br>GTLVTVSA |
| **Variable light chain** | |
| CPS6-IgG | QSALTQPASVSGSPGQSITISCTGTSSDVGTYNHVSWYQQCPGKGPKLLIYDVTNRPSGVSNRFSGSKSDN<br>TASLTISGLQAEDEADYYCYSNTASGAPYVFGSGTKVTVLSQPKANPTVTLFPPSS |
| CPS8-IgG(28H11) | DIVMTQSPSSLAMSVGQKVTMSCKSSQSLLNSSNQKNYLAWYQQKPGQSPKLLVYFASTRESGVPDRFIGS<br>GSGTDFTLTISSVQAEDLADYFCQQHYSTPYTFGGGTKLEIK |
| CPS3-IgG(5F6) | DVVVTQTPLSLPVSLGDQASISCRSSQSLLHSNGNTYLHWYLQK<br>PGQSPKLLIYKVSNRFSGVPDRFSGSGSGTDFTLKISRVEAEDLGVYFCSQSTHVPTF GGGTKLEIK |
| **Constant heavy chain** | |
| IgG1 | ASTKGPSVFPLAPSSKSTSGGTAALGCLVKDYFPEPVTVSWNSGALTSGVHTFPAVLQSSGLYSLSSVVTVP<br>SSSLGTQTYICNVNHKPSNTKVDKKVEPKSCDKTHTCPPCPAPELLGGPSVFLFPPKPKDTLMISRTPEVTCV<br>VVDVSHEDPEVKFNWYVDGVEVHNAKTKPREEQYNSTYRVVSVLTVLHQDWLNGKEYKCKVSNKALPAPIE<br>KTISKAKGQPREPQVYTLPPSREEMTKNQVSLTCLVKGFYPSDIAVEWESNGQPENNYKTTPPVLDSDGSFF<br>LYSKLTVDKSRWQQGNVFSCSVMHEALHNHYTQKSLSLSPGK |
| IgG2 | ASTKGPSVFPLAPCSRSTSESTAALGCLVKDYFPEPVTVSWNSGALTSGVHTFPAVLQSSGLYSLSSVVTVP<br>SSNFGTQTYTCNVDHKPSNTKVDKTVERKCCVECPPCPAPPVAGPSVFLFPPKPKDTLMISRTPEVTCVVVD<br>VSHEDPEVQFNWYVDGVEVHNAKTKPREEQFNSTFRVVSVLTVVHQDWLNGKEYKCKVSNKGLPAPIEKTI<br>SKTKGQPREPQVYTLPPSREEMTKNQVSLTCLVKGFYPSDIAVEWESNGQPENNYKTTPPMLDSDGSFFLY<br>SKLTVDKSRWQQGNVFSCSVMHEALHNHYTQKSLSLSPGK |
| IgG3 | ASTKGPSVFPLAPCSRSTSGGTAALGCLVKDYFPEPVTVSWNSGALTSGVHTFPAVLQSSGLYSLSSVVTVP<br>SSSLGTQTYTCNVNHKPSNTKVDKRVELKTPLGDTTHTCPRCPEPKSCDTPPPCPRCPEPKSCDTPPPCPR<br>CPEPKSCDTPPPCPRCPAPELLGGPSVFLFPPKPKDTLMISRTPEVTCVVVDVSHEDPEVQFKWYVDGVEV<br>HNAKTKPREEQYNSTFRVVSVLTVLHQDWLNGKEYKCKVSNKALPAPIEKTISKTKGQPREPQVYTLPPSRE<br>EMTKNQVSLTCLVKGFYPSDIAVEWESSGQPENNYNTTPPMLDSDGSFFLYSKLTVDKSRWQQGNIFSCSV<br>MHEALHNRFTQKSLSLSPGK |
| **Constant light chain (*kappa* class)** | |
| IgG1,2,3, | RTVAAPSVFIFPPSDEQLKSGTASVVCLLNNFYPREAKVQWKVDNALQSGNSQESVTEQDSKDSTYSLSSTL<br>TLSKADYEKHKVYACEVTHQGLSSPVTKSFNRGEC |
| **Constant light chain (*lambda* class)** | |
| IgG1 | EELQANKATLVCLISDFYPGAVTVAWKADGSPVKAGVETTKPSKQSNNKYAASSYLSLTPEQWKSHRSY<br>SCQVTHEGSTVEKTVAPTECS |
